# Late gestational exposure to flutamide alters stromal composition and immune landscape in the rat mammary gland during pre-puberty, peri-puberty, and adulthood

**DOI:** 10.64898/2026.01.16.699958

**Authors:** David Tovar Parra, Alec McDermott, Jysiane Cardot, Melany Juarez, Fabien Joao, Rhizlane ElOmri, Line Berthiaume, Bhawna Dhawan, Arash Aghigh, Yann Breton, François Légaré, Géraldine Delbès, Martin Pelletier, Étienne Audet-Walsh, Isabelle Plante

**Affiliations:** Institut National đe la Recherche Scientifique (INRS), Centre Armand-Frappier Santé Biotechnologie, Laval, Québec, Canada.; Axe Enđocrinologie-Néphrologie, Centre đe recherche du CHU de Québec-Université Laval, Québec City, Québec, Canada.; Centre Énergie Matériaux Télécom munications, Institut National de la Recherche Scientifique, Varennes, Québec, Canada; Axe Maladies infectieuses et immunitaires, Centre de recherche du CHU de Québec-Université Laval, Québec City, Québec, Canada.; Department of Microbiology-Infectious Diseases and Immunology, Faculty of Medicine, Laval University, Québec, Canada.; Arthritis Research Center, Laval University, Québec, Canada.

**Keywords:** Mammary gland development, anti-androgen, endocrine disruption, androgen receptor, lipidomic, transcriptomic

## Abstract

Perinatal development of the mammary gland is regulated by hormonal signals that influence cell proliferation, extracellular matrix remodeling, immune cell recruitment, and intracellular signaling. While the role of estrogen in mammary gland development is well established, the impact of androgens remains less understood. To address this gap, we inhibited androgen signaling *in utero* using the anti-androgen flutamide (FLU) and investigated the effects on mammary gland development in rats. Using an integrative strategy combining histology, transcriptomics, lipidomics, cytokine profiling, and high-resolution imaging, mammary tissue were analyzed at pre-puberty (postnatal days (PND) 21), peri-puberty (PND46), and adulthood (PND90). FLU exposure induced subtle, yet significant, alterations in mammary morphology and molecular signatures. At PND21, the FLU exposed group exhibited an increased number of adipocytes with reduced size. Transcriptomic analysis revealed differentially expressed genes at PND21 and enrichment in pathways related to androgen response and immune signaling, but minimal changes at later developmental stages. Lipidomic profiling showed transient disruption in long-chain fatty acid composition at early developmental stages. Cytokine profiling revealed a reduced adaptive immune response at PND46 and PND90, and second harmonic generation imaging demonstrated changes in collagen fiber orientation and density across all developmental stages. These data indicate that prenatal androgen signaling is essential for proper stromal development and the establishment of early transcriptional networks in the mammary gland, with only minor long-term effects on glandular architecture in adult nulliparous females.

## 1. Introduction

E Endocrine-disrupting chemicals (EDCs) are exogenous compounds that interfere with sex steroid hormone signaling and homeostasis and have been associated with dysregulations of mammary gland development (Fenton 2006; Szabo and Vandenberg 2021). Many EDCs exhibit estrogenic, anti-estrogenic, androgenic, and/or anti-androgenic activities, ultimately disrupting the endocrine balance (Pan et al. 2023). Sex hormones are critical for the regulation of mammary gland development. Thus, many animal studies have investigated the impact of EDC exposure on altered breast development and breast cancer risk (Fenton 2006). Rodent studies show that *in utero* or neonatal exposure to ECDs can persistently modify mammary gland morphogenesis, altering ductal and lobular architecture and differentiation (Perrot-Applanat et al. 2018; Sapouckey et al. 2018). Exposure to some EDCs may lead to an increased number of carcinogen-sensitive structures in the gland and raise breast cancer susceptibility (Choi et al. 2014; Liao and Dickson 2002).

Normal mammary gland development proceeds through tightly regulated stages. The epithelial mammary tissue starts to develop in a rudimentary tree in both sexes (female and male) around gestational day (GD) 13 (in rodents)(Liao and Dickson 2002; Wansbury et al. 2011). Androgens such as testosterone and dihydrotestosterone (DHT) begin influencing mammary mesenchymal-epithelial dynamics and cause epithelial regression in males between GD14 and GD16 (Hass et al. 2007; Szabo and Vandenberg 2021). In females, postnatal development is primarily driven by rising levels of estrogen, progesterone, prolactin, and other ovarian-pituitary hormones, which stimulate epithelial cell proliferation, ductal elongation, and terminal end bud (TEB) formation at puberty (Brisken and Ataca 2015). Our recent data revealed a pronounced cellular and molecular shift during the transition from pre-puberty to peri-puberty characterized by an expansion of the epithelial area, a reduction in fatty acid concentration, an increase in adipocyte density, and the activation of immune response pathways (Tovar-Parra et al. 2025b). Such coordinated stromal-epithelial remodeling during mammary maturation. Exposure to xenoestrogens, such as bisphenol A (BPA), accelerates ductal elongation and increases epithelial density in mice (Perrot-Applanat et al. 2018). Moreover, our previous research demonstrated that *in utero* exposure to diethylstilbestrol (DES) significantly altered the stroma composition, leading to an increased adipocyte density, reduced polyunsaturated fatty acid levels, changes in immune cell abundance, and diminished chemokine signaling within the tissue (Tovar-Parra et al. 2025a). Together, these findings highlight that endocrine disruption with specify xenoestrogen not only alters epithelial development but also reprograms stromal architecture and immune signaling pathways essential for normal mammary gland morphogenesis.

In contrast, maternal exposure to anti-androgenic compounds can lead to persistent changes in mammary morphology including increased epithelial density and hyperplastic lesions in adult female offspring (Sapouckey et al. 2018), potentially mediated through upregulation of estrogen receptor *α* (ERα) and Wnt/ļ3-catenin signaling (Gao et al. 2014). This reduction in fat deposition improved glucose-lipid homeostasis and mitigated the accelerated lipid accumulation observed in the study participants (Dumesic et al. 2023; Gruessner et al. 2014). Previous studies in female mice exposed to FLU after mid-puberty appear to have minimal effects on the female mammary gland (Peters et al. 2011). Yet, prenatal FLU (2mg/kg/day) exposure in male rats reduces the normal androgen-driven cell death in the rudimentary mammary gland (Fussell et al. 2015; McIntyreBarlow and Foster 2001). In androgen receptor knockout (ARKO) female mice, mammary development is impaired with accelerated ductal branching, an increased number of TEBs, and activated signaling through insulin-like growth factor I (IGF-1), mitogen-activated protein kinase (MAPK), and estrogen receptor (ER) pathways (Gao et al. 2014; Hickey et al. 2012). Suggesting that inhibition of androgen signaling, particularly during development, may disrupt normal mammary patterning through molecular pathways.

Androgen and AR act as counter-regulators to estrogenic stimulation in the mammary gland of both sexes (Dimitrakakis and Bondy 2009). AR is expressed in the epithelium both luminal and myoepithelial cells, as well as in stromal cells as fibroblasts, endothelial, immune cells, and smooth muscle cells (Choi et al. 2014; Li et al. 2010; Tarulli et al. 2019). Through transcriptional activation of key developmental genes via androgen response elements (AREs), AR signaling suppresses epithelial cell proliferation, reduces ductal branching, and limits ductal elongation into the mammary stroma (also known as the fat pad) (Gao et al. 2014; Tarulli et al. 2019). Conversely, AR antagonism abolishes these effects (Gao et al. 2014). Female ARKO mice exhibit accelerated pubertal mammary ductal expansion with increased TEBs (Gao et al. 2014), and adult female rats exposed to FLU show enhanced epithelial proliferation (Dimitrakakis and Bondy 2009). Stage-dependent effects of AR signaling have also been demonstrated. Peter *et al*. showed that DHT exposure from 5 to 12 weeks of age reduced ductal expansion and branching; in contrast, FLU exposure from 12 to 21 weeks increased branching in female mice (Peters et al. 2011). Together, these findings underscore that androgen signaling plays a crucial role in directing mammary epithelial fate during mammary gland development.

Stromal signaling not only regulates adipogenesis and lipid metabolism but also modulates immune infiltration and extracellular matrix (ECM) remodeling, which together provide the structural and biochemical cues that guide epithelial morphogenesis (Liao and Dickson 2002). Thus, disruption of AR signaling in the stroma has the potential to reprogram the mammary microenvironment in ways that are not evident from epithelial changes alone (Tang et al. 2021). Despite these insights, the impact of anti-androgenic disruption, particularly *in utero,* on female mammary gland development, remains poorly characterized. In this context, FLU, a well-characterized nonsteroidal AR antagonist, represents a valuable tool to probe the role of AR signaling. Here, we hypothesize that prenatal AR antagonism disrupts epithelial and stromal programming, leading to alterations in adipocyte organization, immune cell recruitment, and ECM architecture. In this study, we evaluated the effects of FLU exposure (10 mg/kg/day) from GD16 to GD21 on mammary gland development in female rat offspring. We conducted a comprehensive multi-modal analysis at PND21, PND46, and PND90, employing histological and morphometric analyses, bulk RNA sequencing, lipidomic, cytokine profiling, immunofluorescence, and second-harmonic generation (SHG) microscopy. This integrative approach aims to elucidate the role of AR signaling in mammary stromal development and its downstream long-term influence on epithelial and immune components in adulthood.

## Materials and Methods

### 1.1 Animal model and exposure

All animal procedures followed the guidelines outlined by the Canadian Council on Animal Care and were approved by the Institutional Animal Care and Use Committee at the INRS (protocol 2202-02). Rats were housed on a l2Ligth: l2Dark cycle and fed commercial food (Teklad Global 18% protein, Envigo, Madison, WI) and tap water ad libitum. Two virgin 12-20 weeks-old females in proestrus were caged with one 12-20 weeks-old male overnight. The following day, vaginal smears were performed to identify sperm-positive females, and that day was counted as gestational day 0 (GD0). On GD16, dams were randomly assigned to treatment groups. From GD16 to GD21, each dam was weighed daily and gavaged with corn oil (vehicle controls) or 10 mg/kg/day of FLU in corn oil.

FLU (ref F9397) was obtained from Sigma-Aldrich Canada Ltd. (Oakville, Ontario, Canada). The stock solution was dissolved in Dimethyl Sulfoxide (DMSO) (Sigma-Aldrich, Cat. D8418) at 50 mg/ml. For administration, the working solution with a final concentration of 10 mg/ml was prepared by diluting the stock solution 5-fold in corn oil for oral gavage. Equivalent vehicle concentration (4% DMSO) in corn oil (ref C8267) was administered to control animals. Rats were euthanized by CO2 followed by cervical dislocation, and samples were collected at PND21 (pre-puberty), PND46 (peri-puberty), and PND90 (adulthood). For each time point, between five and seven animals were euthanized from different litters (only one female per litter). Two pairs of mammary glands, one from the thoracic region and the other from the inguinal region, were then excised and weighed.

### 1.2 Mammary gland whole mount analysis

One inguinal mammary gland per animal was dissected and mounted on a large microscope slide, following previously described protocols (McDermott et al. 2025; Tovar-Parra et al. 2025b). Samples were fixed overnight at room temperature (RT) in Carnoy’s solution (6:3:1 ratio of 100% ethanol, chloroform, and glacial acetic acid). Following fixation, tissues were rehydrated through a series of ethanol gradients (70% to 0% in water), stained overnight at RT in carmine alum (2% carmine, 5% potassium aluminum sulfate), dehydrated through serial ethanol baths, cleared in xylene, and mounted with Permount (FisherChemical, Ontario, Canada, no. SP15-500).

Whole mounts were imaged with a Canon PowerShot G9x digital camera on a transilluminator (Henning Graphics TR299343) with a measurement scale. Morphological features were analyzed using ImageJ version 1.54 software (https://imagej.net/Fiji/Downloads). Total gland and epithelial areas were quantified by calibrating the image scale, with measurements expressed in cm^2^. Skeletonized images were generated to assess branching complexity, as previously published (Crobeddu et al. 2022; Tovar-Parra et al. 2025b). For each sample (κ=5-7), 3 to 5 representative regions were analyzed using the Sholl analysis plugin in ImageJ-FIJI (StankoEasterling and Fenton 2015; Stanko and Fenton 2017), to determine the number of branch intersections per cm^2^.

### 1.3 *RNA* extraction

Total RNA was extracted from 100 mg of the 4^th^ left inguinal mammary glands using the Aurum total RNA fatty and fibrous tissue kit (Bio-Rad, Ontario, Canada, no. 7326830) according to the manufacturer’s instructions. RNA integrity number (RIN) was assessed using the Agilent 2100 Bioanalyzer instrument and the Agilent RNA 6000 Nano Kit. All samples showed RIN > 8, indicating sufficient quality for sequencing analysis. RNA concentration and purity were measured using a Nanodrop spectrophotometer (Thermo Scientific). Samples were sent to the Genomic Centre of the Centre de recherche du CHU de Québec - Université Faval for RNA-seq analysis using a HiSeq 2500 (125 bp paired-end sequencing), as previously described (Tovar-Parra et al. 2025b).

### 1.4 RNA sequencing data analyses

Raw sequencing reads were assessed for quality using FastQC v0.12.0. Adapters and low-quality bases were trimmed with TrimGalore v0.6.10, and quality was reassessed using FastQC, followed by summary aggregation with MultiQC v1.2. High-quality reads were pseudo-aligned to the Rattus norvegicus Rnor_6.0 transcriptome (https://may2021.archive.ensembl.org/Rattus_norvegicus) using Kallisto vθ.46.1. The dataset was deposited in NCBI under accession number PRJNA1293432. Differential expression analysis was conducted using DESeq2 v1.42.1 in R software. K-means clustering was employed to characterize group-specific gene expression profiles. Genes with Benjamini-Hochberg adjusted p-values (FDR) < 0.05 and absolute log2 fold change > 1.5 were considered differentially expressed. Gene Set Enrichment Analysis (GSEA) was performed to identify pathways modulated across developmental stages (Kanehisa et al. 2025). Functional enrichment analyses were conducted using Gene Ontology (GO) and Kyoto Encyclopedia of Genes and Genomes (KEGG) enrichment tools using the SRplot package (http://www.bioinformatics.com.cn) and R software environment.

### 1.5 Cryosections and histological analysis

Thoracic mammary glands were embedded in OCT Cryomatrix (Fisher Scientific, Cat. 23-730-571) on dry ice and stored at -80°C. Frozen tissues were cryosectioned (5 µm) and fixed in Bouin’s solution (Sigma-Aldrich, Cat. HT10132) overnight at RT. Masson’s Trichrome staining was performed; sections were sequentially treated with Weigert’s iron hematoxylin (10 min), Biebrich scarlet-acid fuchsin (15 min), phosphomolybdic-phosphotungstic acid (20 min), and aniline blue (5 min), followed by washes, 1% acetic acid (5 min), ethanol dehydration (70, 95, and 100%), and xylene clearing before mounting with Permount (Fisher Chemical, Ontario, Canada, No. SP15-500) (Plante Stewart and Eaird 2011). Images were acquired using a Nikon A1R+ microscope and analyzed with ImageJ. Intact adipocyte areas were delineated manually, and overall adipose area and adipocyte number were quantified using the Adiposoft plugin (Galarraga et al. 2012). Tissue composition was further analyzed by segmenting pixel color intensities (hue, saturation, and intensity) using NIS-Element Nikon’s analysis software, distinguishing epithelial (red), collagen (blue), and adipose (white) regions (Crobeddu et al. 2022).

### 1.6 Lipidomic analysis by Gas chromatography/Flame ionization detector

Snap-frozen inguinal mammary glands were pulverized in liquid nitrogen and weighed prior to homogenization in 0.9% saline. The internal standard phosphatidylcholine C2l:O (Sigma-Aldrich) was added before lipid extraction with chloroform: methanol (2:1, v/v), following a modified Folch method (Shaikh and Downar 1981). After centrifugation, the organic phase was collected, dried under nitrogen, resuspended in methanol/benzene (4:1, v/v), and methylated with acetyl chloride as described by Tovar *et al*. (Tovar-Parra et al. 2025b). Fatty acid profiles were analyzed via capillary gas chromatography using an HP5890 gas chromatograph (Hewlett-Packard, Toronto, Canada), an HP-88 capillary column (100 m × 0.25 mm i.d. × 0.20 µm; Agilent Technologies), and a flame ionization detector. Helium was used as the carrier gas (50:1 split ratio). Identification was based on retention times compared to standard mixtures. Data were expressed as % total fatty acids and mg FA/g tissue. Principal component analysis (PCA) was conducted using R, and Pearson correlations were calculated with GraphPad Prism 10.

### 1.7 Immunofluorescence

Thoracic mammary gland cryosections (5 µm) were fixed in 4% formaldehyde, blocked in 3% BSA, and incubated with primary antibodies (Supplementary Table 1) for 60 minutes at RT. Following three 5-minute PBS washes, sections were incubated with the appropriate secondary antibody (Supplementary Table 1) for 60 minutes at RT. Nuclei were counterstained with DAPI (Thermo Scientific, Cat. 62248), and slides were mounted with Fluoromount-G (Cedarlane, Burlington, Ontario, Canada). Images were captured using a Nikon A1R+ confocal microscope. Analysis was performed using Fiji-ImageJ. Negative controls (DAPI-only primary antibody-only and secondary antibody-only) were employed to establish a baseline and fine-tune laser gain, offset, and intensity. For each animal, 4-6 fields were analyzed per gland (n = 5-6 animals per group). Cell quantification was conducted in ImageJ using the “Analyze Particles” tool after converting the image to 8-bit and adjusting the threshold (Image > Adjust > Threshold). To exclude debris and edge cells, size and circularity parameters were optimized. Finally, cell counts were normalized to the total image area to ensure comparable values across samples.

### 1.8 Cytokine and chemokine analysis

The remaining pulverized mammary glands (for RNA extraction) were used for the cytokine and chemokine analysis. Protein extraction was performed using ProcartaPlex™ Cell Tysis Buffer Kit (Cat. N^Q^ EPX-99999-000, Thermo Fisher Scientific), following the manufacturer’s instructions. Protein concentrations were determined with the Pierce BCA kit (Thermo Scientific, Rockford, Illinois, USA), and all samples were normalized to 10 mg/ml. Cytokines and chemokines were quantified using the Cytokine & Chemokine 22-Plex Rat ProcartaPlex™ Panel (Cat. N^Q^ EPX220-30122-901, Thermo Fisher Scientific) on a Luminex platform. Capture beads were added to each well and washed with a magnetic plate washer. Then, 25 µL of assay buffer and 25 µL of sample or standard were added per well. Following a 90-minute incubation and washing, a biotinylated detection antibody mix (25 µL) was added for 30 minutes. After a final wash, 50 µL of Streptavidin-PE was added for 30 minutes. Readings were acquired using a Bio-Plex™ 200 system (Bio-Rad).

### 1.9 Organization of collagen fiber by second harmonic generation microscopy

A second harmonic generation (SHG) microscopy was employed on 5 µm cryosections using a custom-built system equipped with a Ytterbium fiber laser (MPB Communications Inc., Montréal, Canada; 125-fs pulse duration, lO3Onm wavelength, 25 MHz repetition rate). Output power (30-50 mW) was controlled via a half-wave plate and a Glan-Thompson polarizer. A lO× air objective (UPlanSApo, Olympus, Japan; NA 0.3) was used for imaging an area of ~6000 × 6000 µm^2^. Z-axis adjustments were made with piezoelectric motors (PI Nano-Z, USA). SHG and fundamental signals were collected through a condenser, passed through 720 nm short-pass filters and a 515/30 nm band-pass filter (Semrock), and detected with a photomultiplier tube (Hamamatsu R6357, 800 V). Acquisition was synchronized using a multichannel I/O board (National Instruments) and a custom Python program (Pinsard 2020). Images were processed using Fiji-ImageJ. Polarization-SHG (P-SHG) was conducted using linearly polarized light rotated via a motorized half-wave plate (Ducourthial et al. 2019; Golaraei et al. 2020). A series of 18 images was acquired in 20° increments (0°-340°). Collagen fiber orientation (θ) was calculated from intensity variation according to:

I_SHG(Ω) = K [A cos(4Ω − 4θ) + B cos(2Ω − 2θ) + 1],

Where *K* is the mean photon count, and *A* and B are susceptibility constants. Full methodological details can be found in previous publications (Aghigh et al. 2024; Teulon et al. 2015).

### 1.10 Statistical analysis

All data are presented as mean ± standard error of the mean (SEM). Statistical analyses were performed using GraphPad Prism 9.0 (GraphPad Software, San Diego, CA, USA) unless otherwise noted. For all analyses, all samples were coded, and experimenters were blinded to the treatment group during imaging and quantification to prevent any potential bias from the observers. For comparisons between two groups (Control vs FLU) at a single time point, an unpaired two-tailed Student’s ŕ-test was used. When more than two groups or time points were involved, two-way analysis of variance (ANOVA) was applied, followed by Tukey’s post hoc test for multiple comparisons. The assumption of normality was checked with the Shapiro-Wilk test, and data were log-transformed if necessary to meet parametric test assumptions. A *p-* value < 0.05 was considered statistically significant. In figures, significance levels are indicated as *p* < 0.05 (*), *p < 0.01 (**), and p < 0.001* (***). For the transcriptomic data, an adjusted µ-value (FDR) < 0.05 was used to determine significant differential expression, as described above. All statistical tests were two-sided and interpreted conservatively.

## 2. Results

### 2.1 Morphological impact offlutamide after in utero exposure

To assess the impact of *in utero* AR antagonism on postnatal mammary gland development, rat offspring were analyzed at pre-puberty (PND21), peri-puberty (PND46), and adulthood (PND90) (Supplementary Fig. 1A). Body weight and mammary gland weight increased progressively with age in both control and FLU groups without showing significant differences between the two groups (Supplementary Fig. 1B). Mammary gland morphogenic analysis using whole mount showed that epithelial tissue area and epithelial branching followed a similar pattern, increasing with aging but without any significant difference between control and FLU groups (Supplementary Fig. 1C). FLU treatment did not significant impact the epithelial area and the branching at PND21, PND46 and PND90 in FLU-treated animals (Supplementary Fig. 1D-E). These results suggest that *in utero* flutamide exposure does not prevent overall mammary gland ductal development during the transition from early postnatal stages to adulthood.

### 2.2 Transcriptomic patterns reveal disruption of gene expression by Flutamide in a stage-dependent manner

Our previous studies on normal mammary gland development revealed that the transcriptomic profile undergoes significant major transitions from PND21 to PND46 and PND90, with distinct pathway-specific alterations (Tovar-Parra et al. 2025b). To determine whether *in utero* FLU exposure disrupts these developmental transcriptional programs, we conducted bulk RNA-seq on mammary glands collected at PND21, PND46, and PND90. The differential gene expression analysis was performed using p-value < 0.05 and log2FC 1.5. Using these parameters, 414 genes were significantly altered at PND21 (317 downregulated and 97 upregulated), 128 genes were affected at PND46 (79 downregulated and 49 upregulated), and 208 genes were altered (89 downregulated and 119 upregulated) at PND90 (Fig. 1B). We thus transitioned to gene set enrichment analysis (GSEA) to study whether gene signatures, rather than individual genes, would be more globally affected by developmental FLU exposure. Interestingly, the *in utero* exposure to an anti-androgen was associated with an enrichment of the androgen response gene set at all developmental stages, suggesting of a compensatory mechanism later in life (Fig. 1C). Furthermore, an enrichment of estrogen response gene set was also observed at the PND2l and PND9O developmental stages, indicating that both the androgen and estrogen signaling pathways are modulated in the FLU group (Fig. 1C).

**Figure 1.**
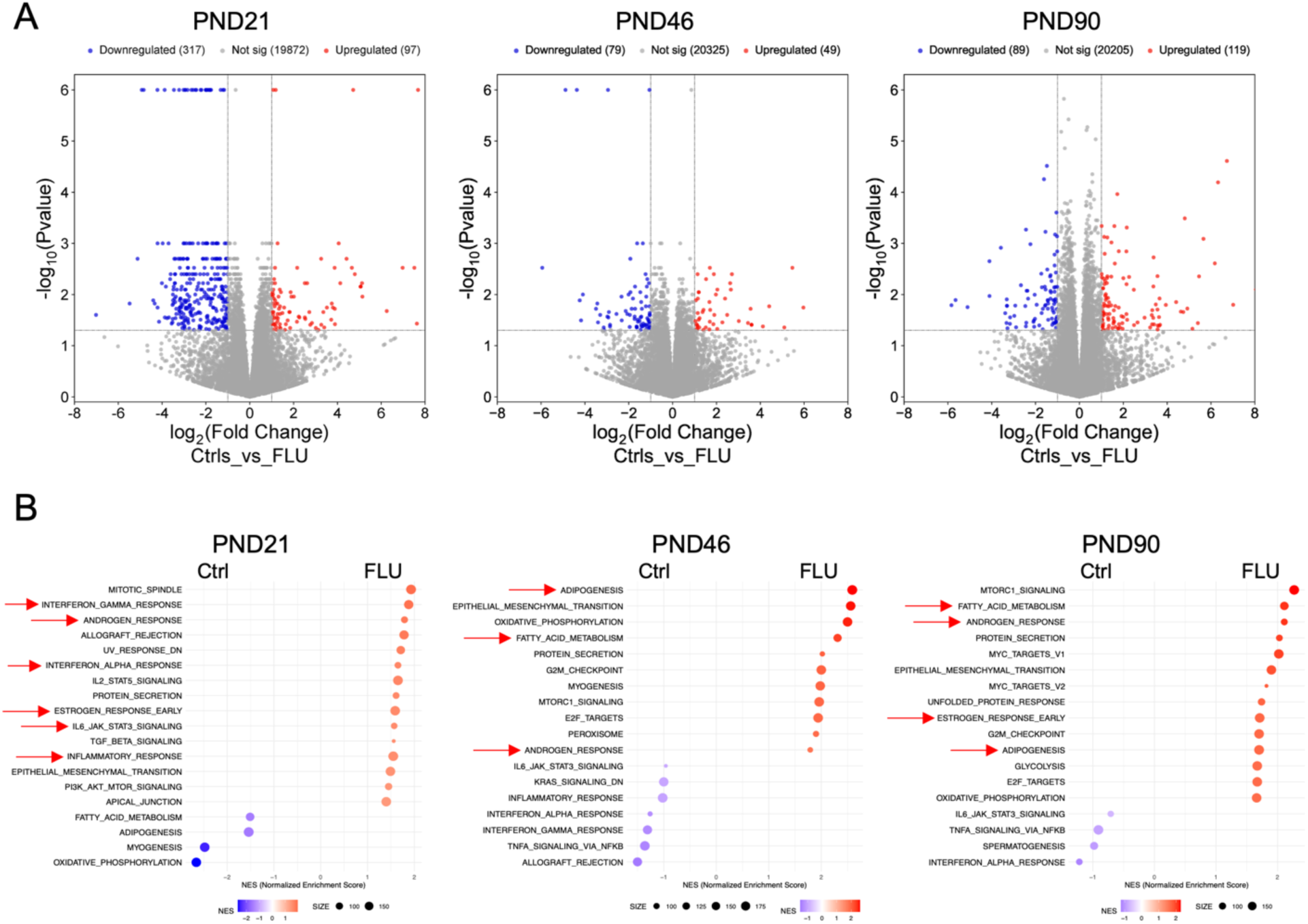
Transcriptional changes in mammary gland development after *in utero* exposure to flutamide. **(A)** Number of differentially expressed genes (DEGs) between flutamide and control groups at each stage, using *p*-value < 0.05 and log_2_FC > 1.5. (**B**) Gene set enrichment analysis (GSEA) of selected biological pathways enriched in the flutamide and control groups at each time point. Data represent n = 4 per group.

During mammary gland development, immune cells are essential to ensure that the epithelium migrates throughout the fat pad and supports alveolar expansion (Reed and Schwertfeger 2010). Resident macrophages, in particular, assemble collagen fibers around TEBs (Vickers and Porter 2024). Interestingly, immune-related response (interferon-α/γ, inflammatory response, IL6-JAK-STAT3 signaling) was enriched at PND21 in FLU-treated animals, whereas control groups revealed a significant enrichment of fatty acid metabolism and adipogenesis signatures (Fig. 1C). In contrast, fatty acid metabolism, adipogenesis, cell cycle control, and epithelial-mesenchymal transition pathways were enriched at PND46 and PND90 in the FLU-treated animals, while immune-related response was enriched in control group (Fig. 1C). Our previous transcriptomic data revealed that immune response pathways are enriched and upregulated at PND46, and maintained until PND90 (Tovar-Parra et al. 2025b). These data suggest that *in utero* exposure to FLU triggers specific developmental stage transcriptional pattern changes in the entire mammary gland micro-environment from pre-puberty to puberty that remain until adulthood stages.

### 2.3 Flutamide alters adipocyte density and tissue composition in the mammary gland

During normal mammary development from pre-puberty to peri-puberty adipocyte density increased while individual size decreased due to stromal remodelation (Tovar-Parra et al. 2025b). To evaluate whether the transcriptomic alterations in adipogenic and metabolic pathways translated into structural changes, we analyzed Masson’s Trichrome-stained sections to quantify adipocyte number, size, and overall tissue composition (Fig. 2A-B). At PND21, the FLU group exhibited significantly increased adipocyte density compared to controls, with a concomitant decrease in mean adipocyte size, suggesting compensatory hyperplasia (Fig. 2C). This increase in adipocyte density persisted through PND46 and PND90, but with no difference in adipocyte size. Tissue segmentation analysis revealed a significant reduction in the percentage of adipose tissue in the FLU group at PND46 and PND90, accompanied by a subtle proportional increase in collagen-rich stromal components (Fig. 2D). These findings indicate that exposure to an androgen receptor antagonist perturbs the establishment of adipose-rich stroma during key windows of mammary development, leading to a persistent shift in tissue architecture that remains throughout puberty and adulthood.

**Figure 2.**
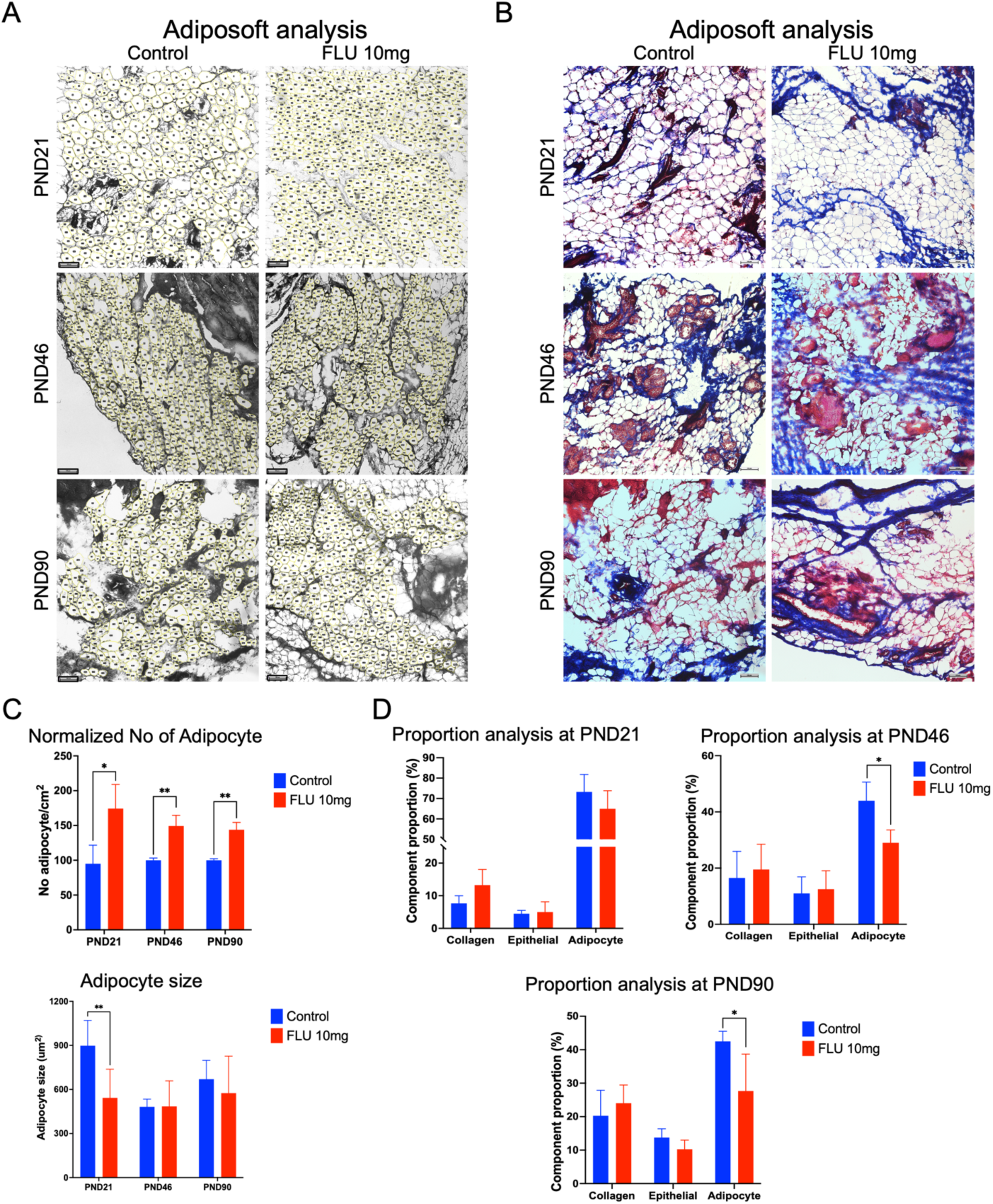
Stromal tissue disruption after *in utero* exposure to flutamide. **(A)** Schematic representation of adipocyte size and number analysis using the Adiposoft plugin from ImageJ at PND21, PND46, and PND90. Scale bar 100 µm. (**B**) Representative Masson’s Trichrome-stained image of a mammary gland section across all developmental stages. Scale bar 100 µm. (**C**) Quantification of adipocyte density and mean adipocyte area. (**D**) Proportional area (%) of mammary gland tissue classified as epithelium, adipose, or collagen-rich stroma based on color segmentation. Bars represent mean ± SD; p < 0.05 (*), p < 0.01 (**).

### 2.4 Flutamide impacts lipidomic profiles in the mammary gland

In our previous studies, we reported that both saturated and polyunsaturated fatty acid concentrations naturally decline from PND21 to PND46 and PND90. To determine whether the transcriptomic pattern of adipogenesis and fatty acid metabolism accompanies alterations in stromal content, we performed lipidomic profiling using gas chromatography on mammary glands. Correlation analysis revealed that FLU disrupted the correlation between fatty acids at PND21, PND46 and PND90 generating a negative correlation, particularly among long-chain polyunsaturated fatty acids (Fig. 3A). Quantitative analysis demonstrated significant reductions in long-chain polyunsaturated fatty acid, including arachidonic acid (20:4 n-6), adrenic acid (22:4 n-6), timnodonic acid (EPA) (20:5 n-3), and docosapentaenoic acid (DPA) (22:5 n-3) in FLU group at PND21 (Fig. 3B). Although FLU also dysregulated the correlation patterns at PND46 and PND90, these differences were not sustained at later stages (PND46 and PND90), supporting the notion that FLU transiently impairs lipid metabolic programming during critical windows of postnatal development.

**Figure 3.**
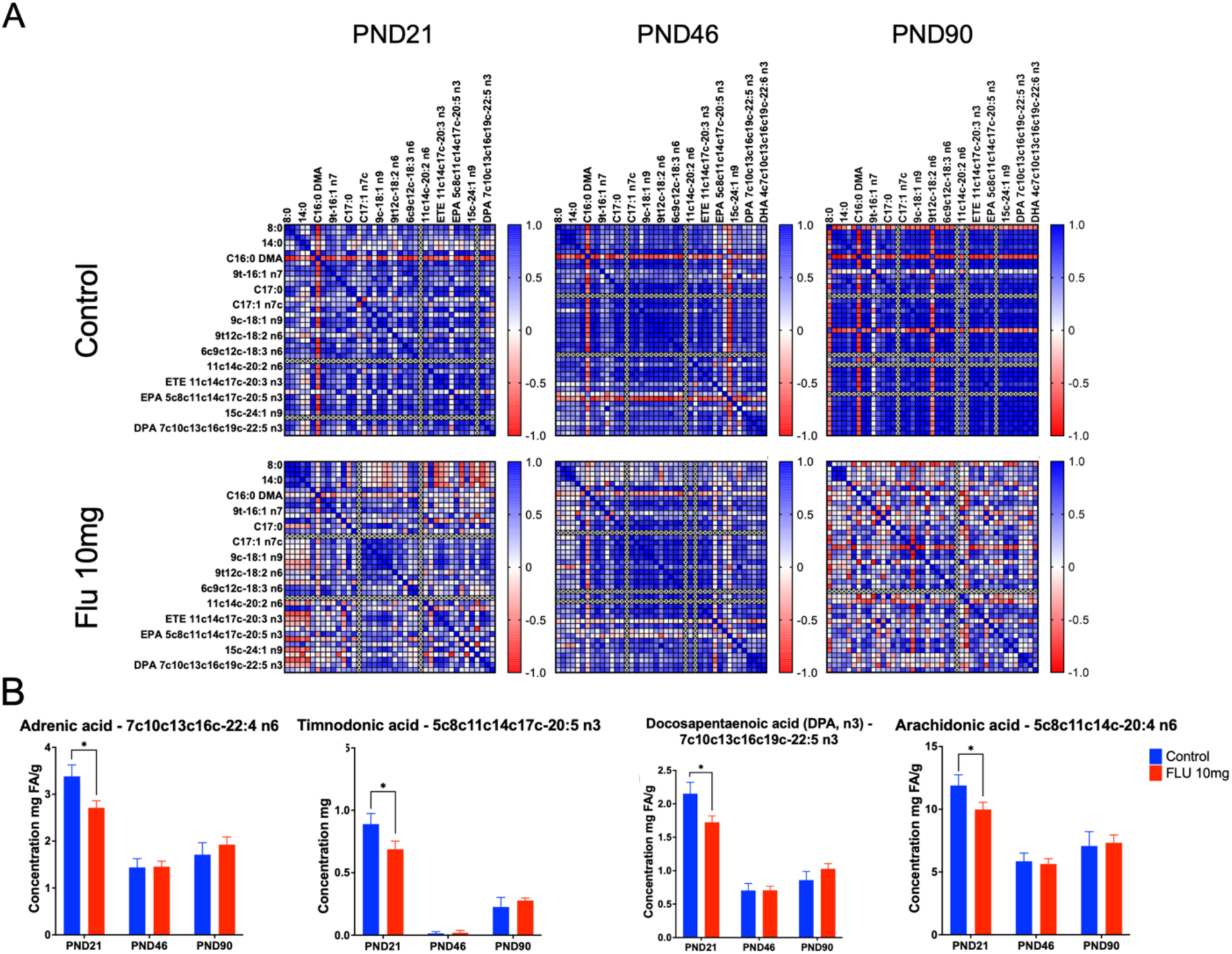
Flutamide alters the mammary gland lipid composition during early postnatal development after *in utero* exposure to flutamide. **(A)** Correlation analysis of faby acid abundance showing pairwise Pearson coefficients within each group; red indicates a negative correlation in the interaction between lipids, and blue indicates a positive correlation. (**B**) Quantification of selected long-chain polyunsaturated faby acids: adrenic acid (22:4 n-6), EPA (20:5 n-3), DPA (22:5 n-3), and arachidonic acid (20:4 n-6). Statistical comparisons performed by unpaired two-tailed t-test; Bars represent mean ± SD; *p* < 0.05 (*).

### 2.5 Blocking the androgen receptor impairs immune cell microenvironment and chemokine signaling during mammary gland development

During the transition from PND21 to PND46, our previous data revealed increased expression of immune-related genes *(Tnfrsflla, Irf7, Traf1,* and *Cd3é)* during peri-puberty (Tovar-Parra et al. 2025b). To investigate whether FLU affected immune dynamics in the mammary gland, we performed cell-type deconvolution from bulk RNA-seq data using M8 cell-type signature gene analysis from GSEA and validated key findings by immunofluorescence and cytokine quantification. As shown in Fig. 4A, transcriptomic deconvolution analysis showed an increase in macrophages, monocytes, and antigen-presenting cells in the FLU group at PND21, but a reduction in lymphocyte-related signatures (T cells, B cells, and NK cells) in the FLU group at PND46 and PND90. Previous studies reported that during pre-puberty, immune cell numbers and expression of key genes *(Cd3e, Cd19, Cd28, Cd69,* and *Il7r)* are normally low (Gopalakrishnan et al. 2018; Reed and Schwertfeger 2010).

**Figure 4.**
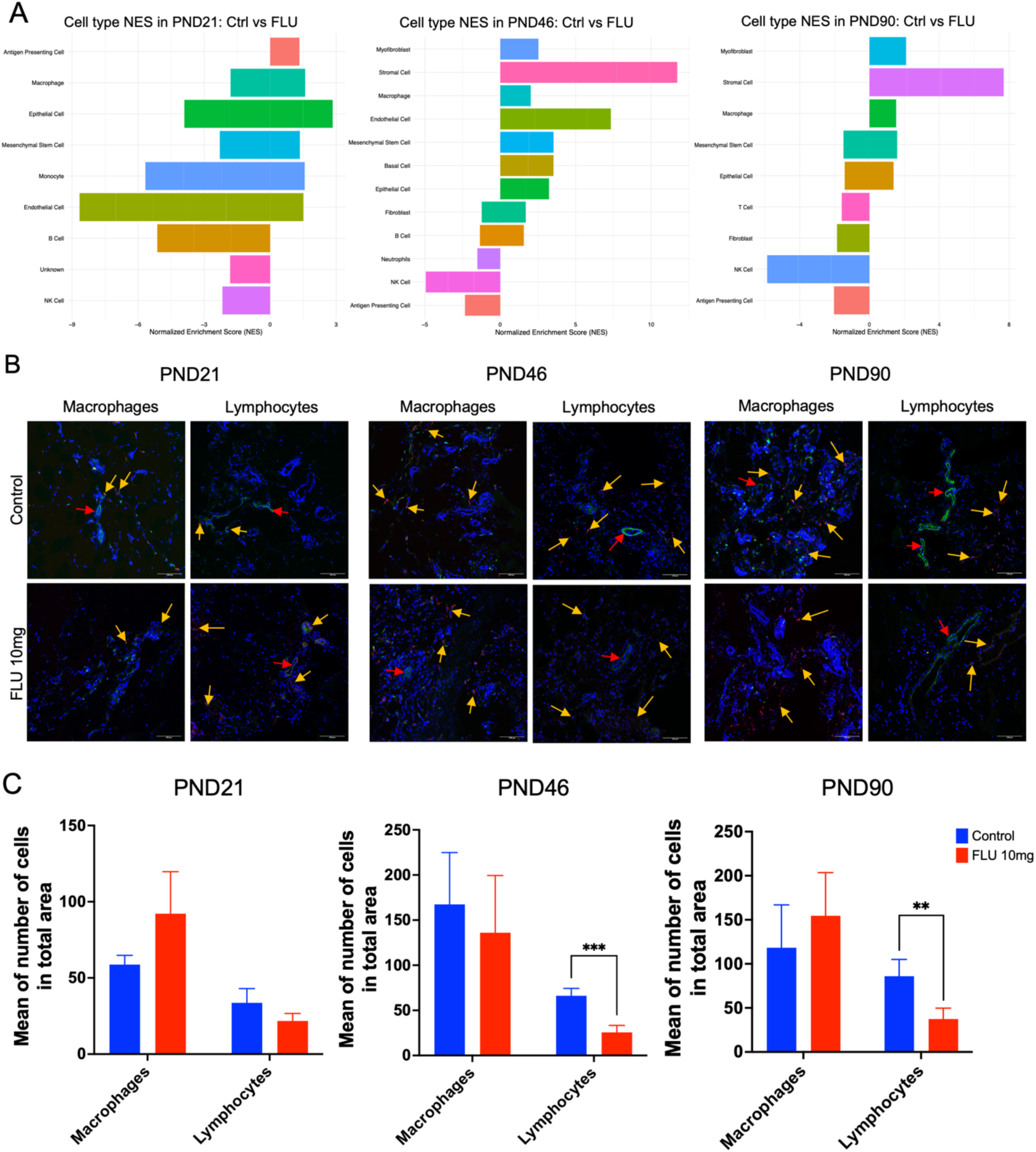
Flutamide impairs lymphocyte recruitment in the mammary gland development. **(A)** Immune cell-type signature from bulk RNA-seq data using deconvolution algorithms from GSEA M8 cell type gene sets. (B) Representative immunofluorescence images of CD45^+^/CD68^+^ Hematopoietic (green)/macrophages (red), CD45^+^/CD3^+^ lymphocytes (red), and DAPI (blue) in mammary tissue at PND2l, PND46, and PND9O. Scale bar 100 µm. Yellow arrow, marking macrophages and lymphocytes, and red arrows, marking hematopoietic cells (blood vessels). **(C)** Quantification of CD45^+^/CD68^+^ and CD45^+^/CD3^+^ cells in total area. Data presented as mean ± SD; p < 0.01 (**), p < 0.001 (***).

We validated these results by immunofluorescence staining, confirming a significant decrease in CD45^+^/CD3^+^ lymphocyte infiltration at PND46 and PND90 stages (Fig. 4B-C). CD45^+^/CD68^+^ macrophage abundance remained unchanged upon FLU treatment at all developmental stages, although total macrophages and lymphocyte numbers were lower at PND21 compared with PND46 and PND90. According to the localization of the immune cell population, we found that CD45^+^ hematopoietic cells and CD68^+^ macrophages are localized around the epithelial tissue and the TEBs. The distribution of lymphocytes (CD45^+^/CD3^+^) is dispersed within the epithelial and stromal tissue (Fig. 4B and Supplementary Fig. 3). Plaks *et al*. reported that CD4^+^ and CD8^+^ cells cluster near ducts and antigen-presenting cells during puberty to promote proper T-cell activation (Plaks et al. 2015). These data suggest that androgen signaling modulates the timing and extent of lymphocyte recruitment during late postnatal gland development.

Given the alteration in the immune cell population caused by FLU, in particular the reduction of lymphocytes at PDN46 and PND90, we then investigated whether FLU could alter the cytokine and chemokine signaling landscape. Previous studies indicate that innate immune-derived TNF cytokine secretion promotes pubertal development, whereas adaptive immune cytokines such as INFγ restrict luminal differentiation (Plaks et al. 2015; Vickers and Porter 2024; Zirbes et al. 2021). PCA revealed different patterns at PND21, where chemokine vectors were moderately associated with the FLU group, and cytokines with the control groups (Fig. 5A). In addition, at PND46, we found a strong variation in the cytokine and chemokine profiles between FLU and control groups, with cytokines and chemokines still more associated with the control group. At PND90, we observe an overlap in the cytokine and chemokine profiles between the FLU and control groups, with high variation in the FLU group.

**Figure 5.**
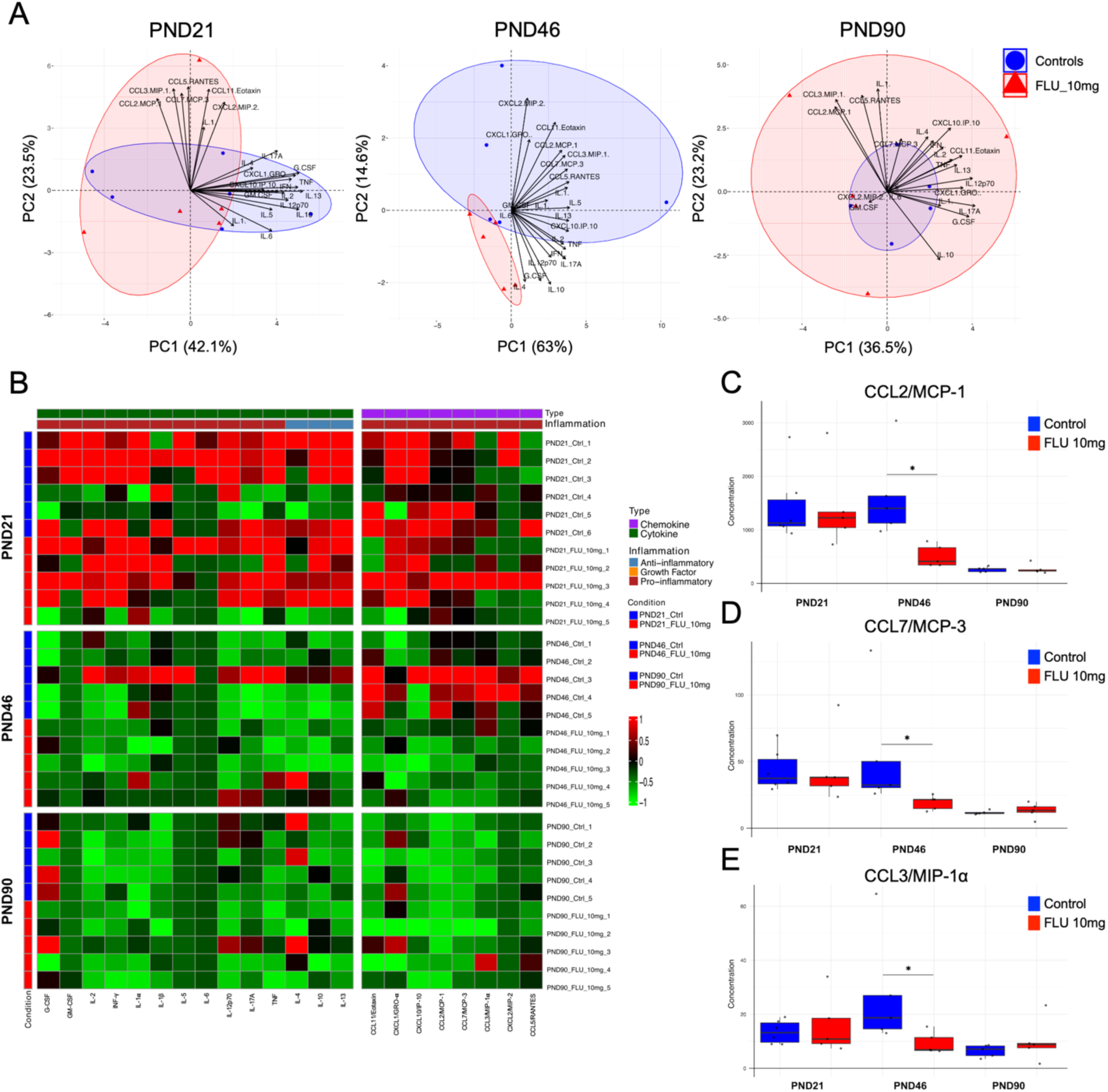
Flutamide reduces chemokine production during peri-pubertal mammary gland development. (**A**) PCA of cytokine and chemokine concentration profiles in mammary tissue lysates at PND21, PND46, and PND90. (**B**) Heatmap showing concentration levels of 22 cytokines and chemokines quantified by multiplex Luminex assay. Individual concentration values (pg/mg protein) for CCL2 (MCP-1) (**C**), CCL7 (MCP-3) (**D**), and CCL3 (MIP-1α) (**E**) at each stage. Bars represent mean ± SD; (*) p < 0.05.

To evaluate and visualize the expression of multiple cytokines and chemokines across all developmental stages, heatmaps were generated, comparing the FLU and control groups at all stages. We found that higher concentrations of cytokines and chemokines were expressed at PND21 compared with PND46 and PND90; however, no differences were found in the patterns of expression between FLU and control (Fig. 5B). Multiplex quantification of cytokines and chemokines further revealed a reduction in pro-inflammatory signals in the FLU group, particularly at PND46. Notably, the levels of CCL2/MCP-1, CCL3/MCP-3, and CCL7/MCP-3, key chemokines involved in leukocyte recruitment, were significantly lower in the FLU group at PND46 (Fig. 5C-E), supporting the observed decrease in lymphocyte infiltration. Teller *et al*. identified Th1 cytokines (IL-2, INFγ) as macrophage activators, while Th2 cytokines (IL-4, IL-5, IL-6, IL-10, IL-13) mediate matrix deposition and tissue remodeling (Reed and Schwertfeger 2010; Teller and White 2009). These results suggest that prenatal FLU exposure reduces the transient inflammatory cues necessary for immune cell microenvironment to the mammary stroma during peri-puberty potentially compromising immune-epithelial crosstalk.

### 2.6 Flutamide exposure alters extracellular matrix composition and collagen remodeling

Given the observed changes in stromal composition, lipidomic profile, and transcriptional patterns, we next examined the extracellular matrix (ECM) by immunofluorescence staining (Collagen I, III, V, and laminin) and second harmonic generation (SHG) microscopy. Quantification of ECM proteins (Fig. 6) showed a significant increase in laminin deposition at PND21 and PND46 in the FLU group (Fig. 6A-B), suggesting altered basal membrane maturation, while collagen I and III levels remained comparable to the control group. In contrast, collagen V puncta were significantly reduced in the FLU group at PND21, suggesting delayed fibrillar collagen deposition (Fig. 6A-B and Supplementary Fig. 4).

**Figure 6.**
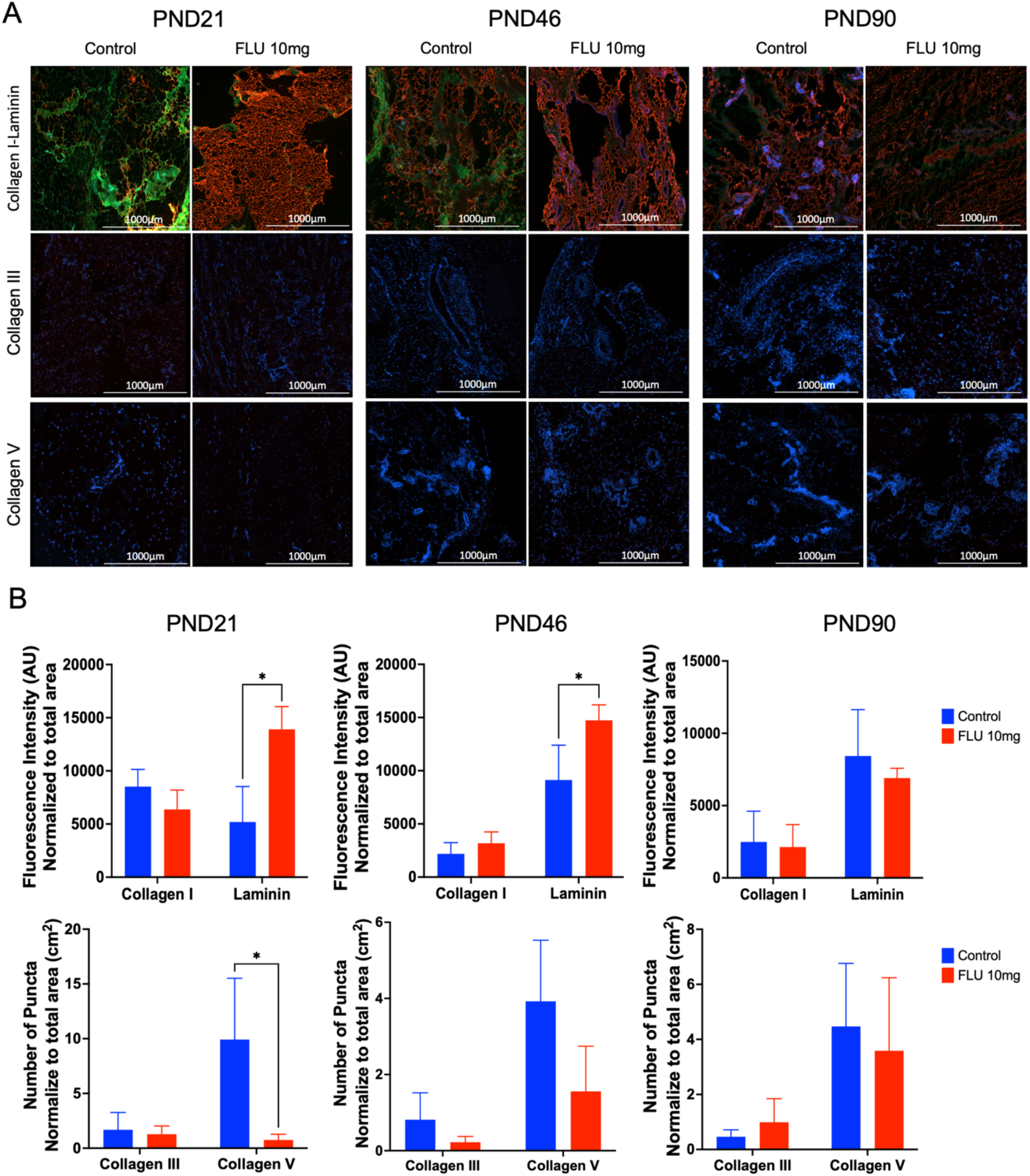
Flutamide modulates ECM protein composition in the mammary gland development. (**A**) Representative immunofluorescence analysis of collagen I (red) and laminin (green) (**A**), Collagen III (red) (**B**), and Collagen V (red) (**C**) across developmental stages. 10x magnification (scale bar = 1000µm). (**D**) Histogram of fluorescence intensity quantifying normalized to total area per cm^2^. Bars represent mean ± SD; (*) p < 0.05.

To further characterize ECM architecture, SHG microscopy was performed (Fig. 7A). Across all developmental stages, the collagen fibers were more aligned to certain angles, but in flutamide exposure resulted in smaller collagen fibers across all time points, with the most pronounced differences at PND46 (Fig. 7B). Orientation histograms confirmed a broader angular distribution in the FLU group, indicating impaired collagen alignment mainly between 30^Q^ a 70^Q^ to -60^Q^ a -20^Q^ (Fig. 7C). Previous studies have showed that properly oriented collagen I fibers facilitate branching morphogenesis and facilitates epithelial cell invasion trhought the fat pad (Brownfield et al. 2013; Schedin and Keely 2011). Together, these data demonstrate that *in utero* exposure to androgen receptor antagonist (FLU) impacts the ECM, causing increased deposition of laminin and a deficient of collagen V with a change in the collagen orientation.

**Figure 7.**
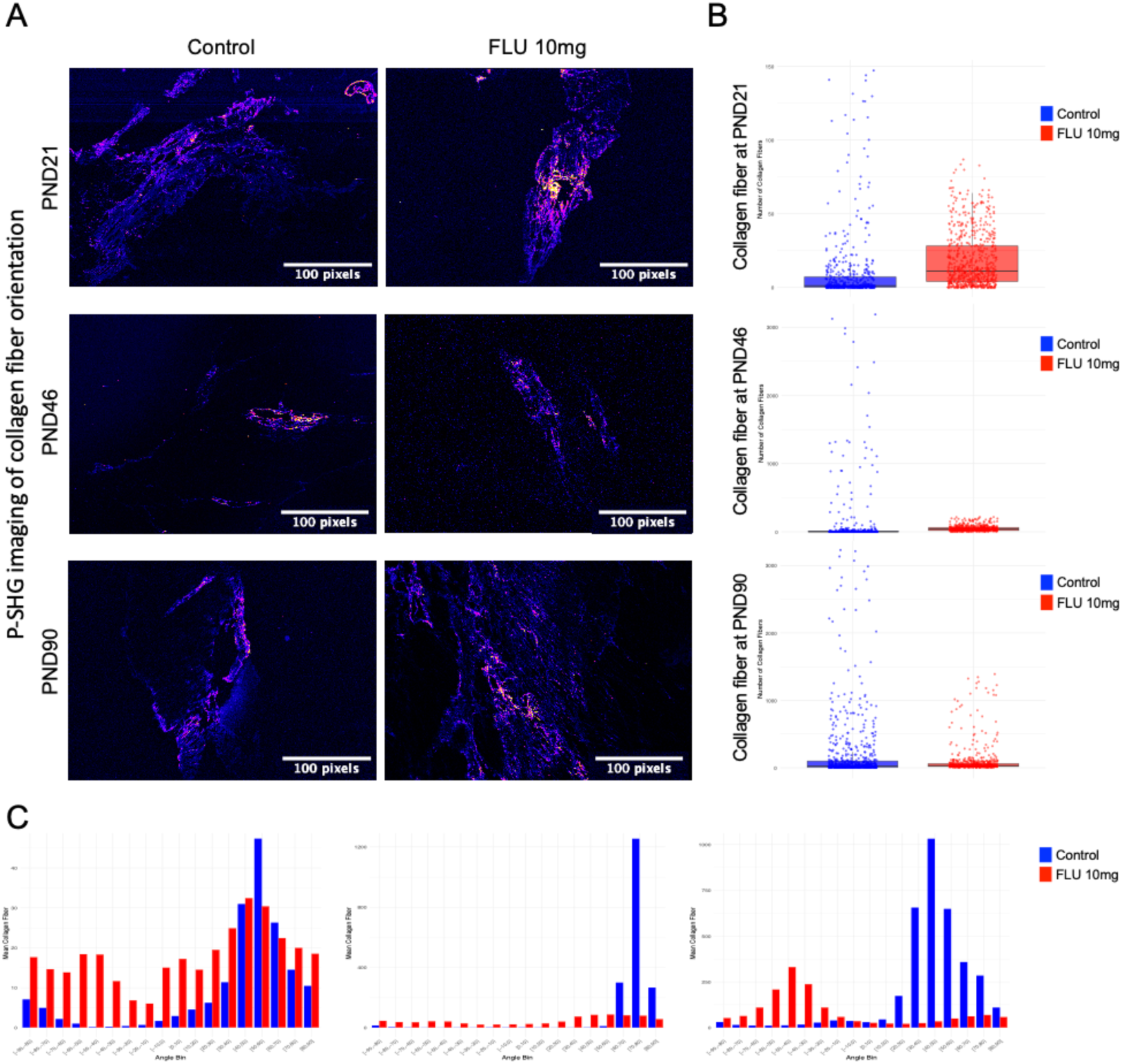
Flutamide disrupts collagen fiber organization and orientation in the mammary gland. (**A**) Representative second harmonic generation (SHG) images of mammary gland stroma at PND21, PND46, and PND90 showing collagen fiber distribution in flutamide and control groups. Scale bar 100 pixels. (**B**) Quantification of collagen fiber density at PND21, PND46, and PND90 in flutamide and control groups. (**C**) Boxplot showing angular distribution of collagen fiber orientation calculated from polarization resolved SHG (p-SHG) across angle bins (from -90° to 90° in 10° intervals) for flutamide and control group.

## 3. Discussion

Postnatal mammary gland development is classically regulated by ovarian hormones, with estrogen and progesterone driving ductal elongation and branching, and prolactin supporting alveologenesis, among others (Sternlicht et al. 2006). However, mammary morphogenesis begins *in utero* and is strongly influenced by mesenchymal-epithelial interaction (Lan et al. 2024; Watson and Khaled 2020). During gestation, androgens play an important role in the sexual dimorphism of the gland (Liao and Dickson 2002). In males, AR activation within the mammary mesenchyme promotes regression of the epithelial buds, while in females, the absence of this signal allows epithelial persistence (Tarulli et al. 2019). Studies in rats and mice have shown that prenatal exposure to flutamide prevents nipple regression in males (McIntyreBarlow and Foster 2001; Tate-Ostroff and Bridges 1988). In females, the role of fetal androgen signaling is less understood, but evidence suggests that stromal AR participates in establishing the architecture of the fat pad and ECM that will later guide ductal invasion (Hickey et al. 2012; Liao and Dickson 2002). Prenatal exposure to FLU (167-300 µg/kg/d) in female mice resulted in significantly smaller ductal trees at PND21, but did not prevent normal pubertal development of the epithelium (Sapouckey et al. 2018). Our findings revealed that *in utero* AR disruption using an AR antagonist during a brief period did not affect epithelial morphology later in life. However, our results showed that FLU induced modest yet significant, long-lasting molecular and stromal alterations in the female mammary gland, without dramatic malformations.

### In utero exposure to FLU alters adipogenesis, immune function and stromal composition at pre-puberty

During normal development, at the pre-pubertal stage (PND21), the mammary gland is small with high plasticity and abundant adipocytes, active immune infiltration at TEB sites, and an ECM in the process of remodeling (Hitchcock et al. 2020; Lue and Radisky 2025; Stewart et al. 2019). In our study, *in utero* FLU exposure significantly increased adipocyte density while reducing mean adipocyte size. Comparable adipogenic responses were previously observed in the DNA-binding dependent (DBD) AR KO male mice, where Rana *et al*. reported increased adipocyte number, size, and adiponectin levels, along with concomitant alterations in lipid metabolism genes such as *Aebp1, Ucp3, Vldlr,* and *Ppara* (Rana et al. 2011). In our results, we did not find dysregulation on these genes, but the adipogenesis pathways were enriched. Our data extend these findings to females, suggesting that blocking AR signaling *in utero* alters adipogenesis in the mammary stroma.

We further detected reduced concentration of long-chain polyunsaturated fatty acids (PUFAs) at pre-puberty, including n-6 fatty acids such as adrenic acid (22:4 n-6), and n-3 fatty acids such as EPA (20:5 n-3) and DPA (22:5 n-3), suggesting disruption of lipid homeostasis. These fatty acids are integral components of cell membranes and precursors to bioactive lipid mediators (HarnackAndersen and Somoza 2009; Schmitz and Ecker 2008). Feng *et al*. demonstrated that treatment to 1000 µg/ml for 24 h with n-3 PUFA *in vitro* using the mammary alveolar cell line (MAC-T) and *in vivo* mouse models positively affects the lipopolysaccharide-induced inflammatory response by increasing the secretion of TNF, FL-6, and IL-lļ3 in mammary alveolar cells through the nuclear transcription factor kappa B (NF-κB) signal pathway (Feng et al. 2021). However, previous studies also reported that n-3 fatty acids can inhibit multiple components of immune cell functions, including those of neutrophils (Rodrigues et al. 2016), macrophages (Allam-Ndoul et al. 2017; Liu et al. 2014), and lymphocytes (GutierrezSvahn and Johansson 2019; WhelanGowdy and Shaikh 2016). Interestingly, gene set enrichment analysis at PND21 revealed activation of androgen-response pathways in the FLU group despite AR *in utero* blockade, suggesting a compensatory reactivation of androgen-regulated transcription. Moreover, immune and interferon pathways were enriched at pre-puberty in the FLU group, consistent with reports that adipocyte hyperplasia modifies cytokine secretion and recruits innate immune cells (MichailidouGomez-Salazar and Alexaki 2022; Moghbeli et al. 2021). Despite these transcriptional changes, no significant alteration in macrophage or lymphocyte distribution was observed during pre-puberty.

The ECM organization was also affected, as the FLU group showed reduced collagen V puncta and elevated laminin deposition. Collagen V is known to regulate fibril assembly of collagen I (Wenstrup et al. 2004), potentially altering the mechanical properties and epithelial-stromal signaling within the mammary gland (Barsky et al. 1982; Berendsen et al. 2006; Breuls et al. 2009; Guo et al. 2022). Similar delays in collagen organization were reported following *in utero* vinclozolin-exposure rat mammary glands by El Sheikh Saad *et al*. (El Sheikh Saad et al. 2013). Together, our results reveal an early stromal disruption characterized by small adipocytes, a pro-inflammatory transcriptional profile, and perturbed ECM distribution, demonstrating that a short prenatal disruption of AR is sufficient to reprogram the stromal tissue before transition to puberty.

### Alterations of the stroma remain are still observable at peri-puberty upon an in utero exposure to FLU

During the peri-pubertal stage (PND46), estrogen and IGF-1 drive rapid epithelial cell proliferation, allowing the ductal elongation and branching formation, while adipocytes hypertrophy to support metabolic demands (Macias and Hinck 2012; Singh and Ali 2023). Normally, AR signaling acts as a counterbalance to estrogenic stimulation, as demonstrated by Gao *et al*. who reported accelerated ductal expansion and increased TEBs in female AR-KO mice via upregulated ERα and Wnt/ļ3-catenin signaling (Gao et al. 2014). In our study, the *in utero* FLU group displayed normal epithelial expansion at PND46, consistent with Peters *et al*. who showed that peri-pubertal FLU or DHT exposure had a limited impact on ductal morphology (Peters et al. 2011). Despite these morphological normalizations, stromal differences persisted in our study. An increase in adipocyte number remained evident, consistent with long-term reprogramming of the fat pad cellularity. Transcriptomic data revealed enrichment of adipogenesis and lipid metabolism pathways in the FLU group, paralleling findings by El Sheikh Saad *et al*. who identified 56 and 122 differentially expressed genes related to morphogenesis *(Actn2, Casq1, Ckm, Phil, Mylk2, Pvalb, Pygm,* and *Tnni2f* lipid metabolism *(Actn2, Csn2, Fabp3, Tnni2,* and *Wap),* and development *(Cldn1, Elf5, Krtl7, Sprrla,* and *Tfap2c)* at PND35 and PND50 (peri-puberty stages) using Wistar rat models exposed *in utero* to vinclozolin, a fungicide that competitively antagonizes the binding of natural androgens to their receptor (El Sheikh Saad et al. 2013).

We also found reduced adaptive immune signatures and a significant decrease in lymphocyte number in the FLU group compared to the control group at PND46. Concentrations of key chemokines, including CCL2/MCP-1, CCL3/MIP-1α, and CCL7/MCP-3, were lower in the FLU groups during peri-puberty indicating impaired immune recruitment. Androgens have been associated with exerting anti-inflammatory effects on the immune system (Ainslie et al. 2024). On one hand, innate immune response with AR activation in myeloid cells tends to dampen pro-inflammatory cytokine production and macrophage activation (Becerra-DiazSong and Heller 2020). In macrophages, testosterone/AR signaling increases IL-10 and reduces TNF and IL-l(3, thereby skewing toward a less inflammatory state (Ainslie et al. 2024). On the other hand, other results suggest that AR activation modulates the expression of pro-inflammatory molecules such as TNF in mice (Lai et al. 2009). In addition, adaptive immune responses decreased in AR deficiency of CD4 CD8^+^ T cells (Olsen et al. 1991). Additionally, changes in gene expression were observed in pathways involved in interferon signaling and Th1 differentiation (Kissick et al. 2014). Interestingly, Fijak *et al*. reported that FLU treatment decreased the production of IL-10 in an *in vitro* model (Fijak et al. 2015), supporting an anti-androgenic pro-inflammatory shift. Finally, the ECM architecture also remained altered at PND46. SHG microscopy revealed misaligned collagen fibers and elevated laminin deposition in the FLU group by immunofluorescence. Ingman *et al*. showed that macrophages align collagen fibrils around TEBs to facilitate ductal invasion; thus, the reduced lymphocyte and altered macrophage-ECM interaction in the FLU group may contribute to abnormal collagen orientation (Ingman et al. 2006). These findings suggest that even when epithelial development appears normal, stromal-immune-ECM interactions are compromised by *in utero* AR disruption.

### In utero exposure to FLU induces long-term changes in the mammary stroma

By the adulthood stage (PND90), the mammary gland has a mature epithelial network supported by a relatively quiescent stroma (Biswas et al. 2022; Sternlicht et al. 2006). In our study, FLU and control glands were indistinguishable in overall epithelial organization, consistent with Dimitrakakis *et al*. who found that adult female rats exposed to FLU showed increased epithelial proliferation but not major structural malformations (Dimitrakakis et al. 2003). Similarly, Peters *et al*. reported that AR blockage during mid-puberty (5 weeks) and post-puberty (12 weeks) increased epithelial cell proliferation without altering ER levels (Peters et al. 2011). However, stromal differences persisted in our model. Adipocyte number remained elevated, indicating permanent reprogramming of fat pad cellularity even though the adipocyte proportion is significantly smaller from PND21 to PND46 and PND90. Also, these changes remained in the stroma with the lymphocyte deficit observed at PND46 enduring at PND90. According to Stewart *et al*. immune cell numbers and localization decline during adulthood, concentrating around ducts, side buds, and often between the luminal-basal interface (Stewart et al. 2019). Also, the expression of AR has been negatively correlated with overall immune composition in breast tissue (Ben-Batalla et al. 2020; Hanamura et al. 2023).

Collagen fibers remained less aligned, indicating a lasting ECM signature of early AR disruption. Importantly, these changes were confined to the stroma rather than the epithelium, distinguishing our model from AR KO mice models in other studies, where epithelial growth pathways (ERα, IGF-1, MAPK) are permanently altered (Yeh et al. 2003). These changes in the stroma support the idea that a transient fetal anti-androgenic insult reprograms the stromal microenvironment rather than directly impairing epithelial lineages. In conclusion, our data suggest that *in utero* antagonism of androgen receptors creates a long-lasting stromal disruption that extends beyond adipocyte cellularity, influences chemokine signaling, and alters the extracellular matrix architecture. This subtle disruption subtly guides postnatal mammary development without causing overt epithelial malformations.

## 4. Conclusion

In conclusion, transient late gestation exposure to AR inhibition by FLU resulted in subtle but coherent stromal reprogramming across developmental stages from pre-puberty to adulthood without causing overt epithelial malformations. The effects were most pronounced before and around peri-puberty affecting adipogenesis, lipid metabolism, fatty acid profiles, immune cells and signaling activity and ECM organization. By adulthood, the gland appeared largely normal. Yet, the stromal compartment changes after androgen receptor disruption can modulate the mammary microenvironment. These findings gain particular relevance within the broader context of EDCs that act as antiandrogens via AR antagonism, such as flutamide, vinclozolin, and other anti-androgenic compounds that interfere with hormonal signaling during critical developmental windows. Our results provide insight into how such EDCs exposures can durably reprogram the stromal compartment, potentially predisposing the mammary gland to altered responses later in life. This study emphasizes that the mammary stroma, rather than the epithelium, is a sensitive target of anti-androgenic exposures during development, reinforcing the importance of including stromal endpoints in risk assessments of EDCs. Altogether, our results highlight the concept of early stromal programming, in which endocrine disruption during late gestation leaves latent imprints on tissue architecture that may influence later functional responses to hormonal cycles, pregnancy, or carcinogenic insults.

## Table legends

**Table 1.** Fatty acid profiles in mammary glands across developmental stages after in utero FLU exposure conditions.

| Fatty acids | Carboxyl nomenclature | PND21 (mg FA/g) |  | PND46 (mg FA/g) |  | PND90 (mg FA/g) |  |
| --- | --- | --- | --- | --- | --- | --- | --- |
|  |  | Control | FLU 10mg | Control | FLU 10mg | Control | FLU 10mg |
| Caprylic acid | C8:0 | 1.55 ±0.50 | 1.66 ±0.74 | 0.19 ±0.09 | 0.19 ±0.04 | 0.07 ±0.05 | 0.09 ±0.05 |
| Capric acid | C10:0 | 20.79 ±4.29 | 20.00 ±5.92 | 3.00 ±1.78 | 3.30 ±0.74 | 0.72 ±0.50 | 1.10 ±0.46 |
| Lauric acid | C12:0 | 43.25 ±6.49 | 42.13 ±12.12 | 8.91 ±5.27 | 9.19 ±2.10 | 2.59 ±1.28 | 3.88 ±1.87 |
| Myristic acid | C14:0 | 48.8 ±9.81 | 41.72 ±21.19 | 13.26 ±7.20 | 12.64 ±2.68 | 6.48 ±2.38 | 8.16 ±2.90 |
| Myristoleic acid | 9c-14:1 n5 | 0.75 ±0.48 | 0.96 ±0.62 | 0.50 ±0.21 | 0.38 ±0.13 | 0.38 ±0.13 | 0.32 ±0.32 |
| Pentadecanoic acid | C15:0 | 0.94 ±0.28 | 0.94 ±0.15 | 0.56 ±0.18 | 0.61 ±0.11 | 0.66 ±0.28 | 0.85 ±0.25 |
| 1,1-Dimethoxyhexadecane | C16:0 DMA | 0.02 ±0.04 | 0.02 ±0.03 | 0.07 ±0.05 | 0.07 ±0.05 | 0.06 ±0.04 | 0.02 ±0.03 |
| Pentadecenoic acid (trans isomer) | 10t-15:1 n5 | 0.13 ±0.03 | 0.13 ±0.02 | 0.07 ±0.04 | 0.08 ±0.03 | 0.13 ±0.05 | 0.13 ±0.07 |
| Palmitic acid | C16:0 | 132.00 ±26.26 | 127.50 ±29.32 | 69.72 ±26.24 | 68.49 ±13.04 | 72.51 ±27.60 | 84.62 ±16.49 |
| Palmitelaidic acid | 9t-16:1 n7 | 0.08 ±0.06 | 0.11 ±0.03 | 0.04 ±0.03 | 0.04 ±0.04 | 0.03 ±0.04 | 0.03 ±0.05 |
| Isoheptadecanoic acid | C17:0 isobranched | 0.10 ±0.07 | 0.11 ±0.05 | 0.06 ±0.05 | 0.07 ±0.04 | 0.12 ±0.09 | 0.10 ±0.11 |
| Palmitoleic acid | 9c-16:1 n7 | 12.02 ±4.46 | 13.69 ±5.94 | 11.84 ±5.31 | 9.95 ±4.10 | 12.67 ±4.31 | 13.58 ±8.63 |
| Margaric acid | C17:0 | 0.94 ±0.34 | 0.88 ±0.24 | 0.58 ±0.20 | 0.64 ±0.11 | 0.71 ±0.30 | 0.57 ±0.54 |
| Heptadecenoic acid (trans isomer) | 10t-17:1 n7 | 0.27 ±0.09 | 0.25 ±0.05 | 0.21 ±0.06 | 0.22 ±0.06 | 0.25 ±0.07 | 0.30 ±0.04 |
| 1,1-Dimethoxyoctadecane | C18:1 DMA | 0.12 ±0.31 | 0.00 ±0.00 | 0.00 ±0.00 | 0.00 ±0.00 | 0.00 ±0.00 | 0.01 ±0.02 |
| Heptadecenoic acid (cis isomer) | C17:1 n7c | 0.50 ±0.34 | 0.68 ±0.21 | 0.44 ±0.16 | 0.45 ±0.11 | 0.47 ±0.14 | 0.56 ±0.14 |
| Stearic acid | C18:0 | 17.11 ±4.74 | 16.10 ±3.14 | 10.78 ±3.63 | 11.02 ±1.95 | 12.13 ±4.57 | 13.95 ±3.09 |
| Trans-vaccenic acid | 11t-18:1 n7 | 0.36 ±0.27 | 0.47 ±0.21 | 0.41 ±0.19 | 0.39 ±0.11 | 0.36 ±0.31 | 0.20 ±0.16 |
| Oleic acid | 9c-18:1 n9 | 102.51 ±33.93 | 97.91 ±20.14 | 76.08 ±26.69 | 77.26 ±16.19 | 84.49 ±30.74 | 100.31 ±17.28 |
| Vaccenic acid | 11c-18:1 n7 | 8.25 ±2.67 | 8.17 ±1.64 | 6.35 ±2.09 | 6.77 ±1.27 | 8.30 ±3.12 | 9.41 ±1.41 |
| Petroselinic acid | 13c-18:1 n5 | 0.21 ±0.13 | 0.25 ±0.06 | 0.14 ±0.07 | 0.15 ±0.03 | 0.19 ±0.07 | 0.13 ±0.10 |
| Conjugated linoleic acid (trans, cis isomer) | 9t12c-18:2 n6 | 0.18 ±0.20 | 0.16 ±0.20 | 0.21 ±0.10 | 0.20 ±0.13 | 0.17 ±0.11 | 0.30 ±0.17 |
| Linoleic acid | LA 9c12c-18:2 n6 | 165.54 ±49.82 | 161.30 ±25.36 | 123.50 ±42.60 | 132.30 ±28.77 | 147.50 ±60.59 | 179.20 ±31.12 |
| Arachidic acid | C20:0 | 0.20 ±0.12 | 0.19 ±0.10 | 0.18 ±0.09 | 0.21 ±0.03 | 0.21 ±0.06 | 0.24 ±0.08 |
| Gamma-linolenic acid (GLA) | 6c9c12c-18:3 n6 | 1.93 ±0.65 | 1.90 ±0.40 | 0.74 ±0.32 | 0.73 ±0.17 | 0.70 ±0.33 | 0.81 ±0.11 |
| Alpha-linolenic acid (ALA) | ALA 9c12c15c-18:3 n3 | 12.46 ±3.49 | 11.23 ±3.62 | 7.72 ±2.83 | 8.59 ±2.19 | 9.81 ±4.55 | 11.79 ±2.43 |
| Gondoic acid | 8c-20:1 n12 | 0.00 ±0.00 | 2.44 ±7.34 | 0.00 ±0.00 | 0.00 ±0.00 | 0.00 ±0.00 | 0.00 ±0.00 |
| 11-Eicosenoic acid | 11c-20:1 n9 | 0.87 ±0.55 | 1.26 ±1.00 | 0.76 ±0.28 | 0.80 ±0.15 | 0.59 ±0.39 | 0.58 ±0.46 |
| Stearidonic acid | 6c9c12c15c-18:4 n3 | 0.30 ±0.13 | 0.26 ±0.07 | 0.014 ±0.03 | 0.00 ±0.00 | 0.00 ±0.00 | 0.07 ±0.11 |
| Eicosadienoic acid | 11c14c-20:2 n6 | 3.98 ±1.13 | 3.62 ±1.04 | 1.60 ±0.53 | 1.76 ±0.36 | 1.35 ±0.55 | 1.63 ±0.31 |
| Behenic acid | C22:0 | 0.22 ±0.06 | 0.22 ±0.064 | 0.19 ±0.06 | 0.19 ±0.04 | 0.22 ±0.09 | 0.25 ±0.05 |
| Dihomo-gamma-linolenic acid (DGLA) | 8c11c14c-20:3 n6 | 2.96 ±0.76 | 2.71 ±0.56 | 1.01 ±0.37 | 1.04 ±0.20 | 1.00 ±0.40 | 1.13 ±0.24 |
| Eicosatrienoic acid | ETE 11c14c17c-20:3 n3 | 0.26 ±0.15 | 0.27 ±0.12 | 0.06 ±0.06 | 0.07 ±0.06 | 0.07 ±0.07 | 0.06 ±0.06 |
| Arachidonic acid | AA 5c8c11c14c-20:4 n6 | 11.94 ±2.47 | 10.16 ±1.60 * | 5.85 ±1.72 | 5.64 ±1.17 | 7.07 ±2.53 | 7.33 ±1.52 |
| Docosadienoic acid | 13c16c-22:2 n6 | 0.14 ±0.03 | 0.11 ±0.05 | 0.03 ±0.04 | 0.02 ±0.03 | 0.00 ±0.00 | 0.01 ±0.02 |
| Timnodonic acid (EPA) | EPA 5c8c11c14c17c-20:5 n3 | 0.89 ±0.24 | 0.63 ±0.29 * | 0.01 ±0.03 | 0.01 ±0.05 | 0.22 ±0.17 | 0.27 ±0.05 |
| Lignoceric acid | C24:0 | 0.51 ±0.16 | 0.49 ±0.12 | 0.33 ±0.15 | 0.38 ±0.06 | 0.38 ±0.12 | 0.44 ±0.10 |
| Clupanodonic acid | 13c16c19c-22:3 n3 | 0.00 ±0.00 | 0.01 ±0.04 | 0.06 ±0.04 | 0.06 ±0.04 | 0.10 ±0.04 | 0.08 ±0.07 |
| Nervonic acid | 15c-24:1 n9 | 0.19 ±0.05 | 0.19 ±0.03 | 0.15 ±0.03 | 0.16 ±0.02 | 0.15 ±0.04 | 0.17 ±0.02 |
| Adrenic acid | 7c10c13c16c-22:4 n6 | 3.41 ±0.70 | 2.76 ±0.42 * | 1.43 ±0.48 | 1.45 ±0.33 | 1.71 ±0.57 | 1.92 ±0.40 |
| Docosapentaenoic acid (DPA, n6) | 4c7c10c13c16c-22:5 n6 | 1.01 ±0.20 | 0.93 ±0.26 | 0.50 ±0.16 | 0.47 ±0.10 | 0.56 ±0.25 | 0.55 ±0.16 |
| Docosapentaenoic acid (DPA, n3) | DPA 7c10c13c16c19c-22:5 n3 | 2.20 ±0.46 | 1.74 ±0.25 * | 0.70 ±0.27 | 0.70 ±0.17 | 0.86 ±0.28 | 1.02 ±0.18 |
| Docosahexaenoic acid (DHA) | DHA 4c7c10c13c16c19c-22:6 n3 | 2.76 ±0.58 | 2.52 ±0.59 | 1.16 ±0.36 | 1.05 ±0.23 | 1.43 ±0.71 | 1.46 ±0.22 |
| **Categories** |  |  |  |  |  |  |  |
| Ratio of omega-3 to omega-6 fatty acids | n-3/n-6 | 0.10 ±0.004 | 0.09 ±0.01 | 0.07 ±0.01 | 0.07 ±0.01 | 0.07 ±0.01 | 0.07 ±0.01 |
| Total omega-3 fatty acids | n-6 total | 191.21 ±55.49 | 183.70 ±28.66 | 134.90 ±45.77 | 143.60 ±30.76 | 160.10 ±65.12 | 192.90 ±32.09 |
| Total omega-6 fatty acids | n-3 total | 19.12 ±4.84 | 17.71 ±3.07 | 10.05 ±3.88 | 10.52 ±2.54 | 12.58 ±5.77 | 14.95 ±2.57 |
| Saturated fatty acids | Saturated | 266.32 ±46.59 | 251.90 ±63.73 | 107.80 ±42.42 | 106.90 ±17.93 | 96.71 ±36.81 | 114.20 ±21.57 |
| Ratio of polyunsaturated to saturated fatty acids | P (n-3 & 6)/S | 0.78 ±0.18 | 0.83 ±0.16 | 1.37 ±0.23 | 1.43 ±0.14 | 1.77 ±0.13 | 1.84 ±0.23 |
| Cis polyunsaturated omega-3 fatty acids | Polyunsaturated n3 cis | 19.11 ±4.84 | 17.70 ±3.06 | 10.05 ±3.88 | 10.52 ±2.54 | 12.52 ±5.78 | 14.80 ±2.68 |
| Cis polyunsaturated omega-6 fatty acids | Polyunsaturated n6 cis | 190.80 ±55.43 | 183.40 ±28.59 | 134.60 ±45.71 | 143.40 ±30.67 | 160.00 ±65.19 | 192.61 ±31.96 |
| Trans polyunsaturated omega-6 fatty acids | Polyunsaturated n6 trans | 0.26 ±0.16 | 0.19 ±0.19 | 0.22 ±0.07 | 0.20 ±0.13 | 0.17 ±0.11 | 0.30 ±0.17 |
| Trans monounsaturated fatty acids | Trans monounsaturated | 0.92 ±0.35 | 1.07 ±0.23 | 0.74 ±0.30 | 0.75 ±0.24 | 0.78 ±0.40 | 0.69 ±0.17 |
| Cis monounsaturated fatty acids | Cis monounsaturated | 125.39 ±40.15 | 122.80 ±22.88 | 96.30 ±33.61 | 95.97 ±21.18 | 107.30 ±38.15 | 125.40 ±21.77 |
| Cis monounsaturated omega-6 fatty acids | Monounsaturated n6 cis | 0.00 ±0.00 | 0.02 ±0.06 | 0.00 ±0.00 | 0.00 ±0.00 | 0.00 ±0.00 | 0.00 ±0.00 |
| Total trans fatty acids | Total trans | 1.20 ±0.40 | 1.27 ±0.37 | 0.97 ±0.36 | 0.95 ±0.35 | 1.01 ±0.35 | 1.15 ±0.56 |
| Total cis fatty acids | Total cis | 335.34 ±98.85 | 324.00 ±52.13 | 241.00 ±81.92 | 249.90 ±52.44 | 279.70 ±108.60 | 332.80 ±53.49 |
| Total cis omega-6 fatty acids | Total omega-6 cis | 190.91 ±55.49 | 183.50 ±28.64 | 134.70 ±45.69 | 143.40 ±30.67 | 160.00 ±65.19 | 192.60 ±31.96 |
| Total Fatty acids | Total Fatty acids | 603.00 ±136.71 | 579.10 ±100.80 | 349.90 ±122.90 | 357.90 ±69.17 | 377.70 ±145.60 | 448.30 ±73.44 |

Significance codes: 0.05 \*\*

**Table 2.**
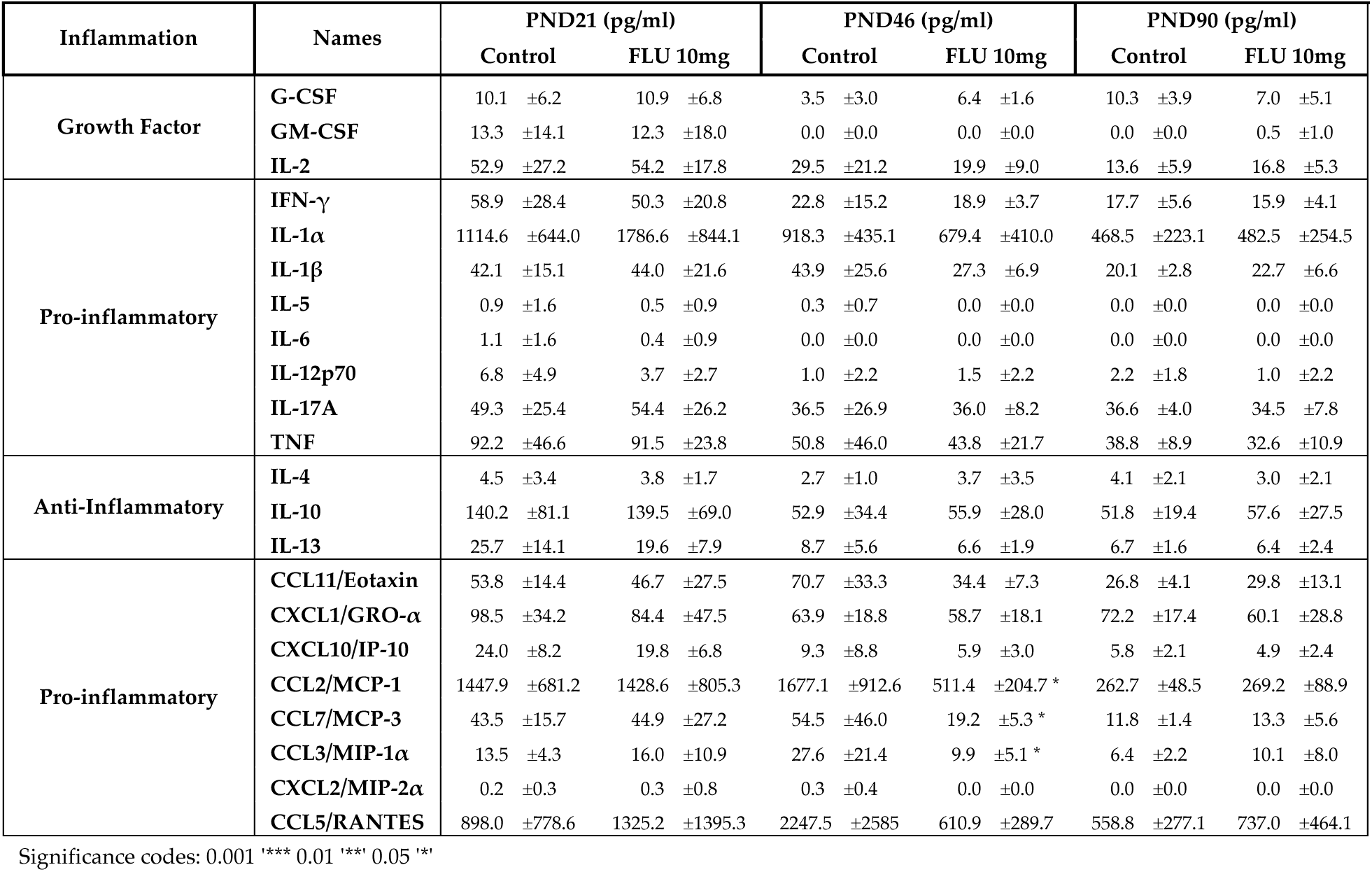
Cytokine and chemokine profiles across developmental stages after in utero FLU exposure.

## Supplementary material

**Supplementary Table 1.** List of Antibodies used for immunofluorescence.

| Target protein | Host | Dilution | Manufacturer | Catalogue # |
| --- | --- | --- | --- | --- |
| **Primary antibodies** |  |  |  |  |
| Collagen I | Mouse | 1:400 | Invitrogen | MA1-26771 |
| Collagen III | Rabbit | 1:200 | Invitrogen | PA5-99160 |
| Collagen V | Rabbit | 1:400 | Invitrogen | PA5-102416 |
| Laminin | Rabbit | 1:400 | Invitrogen | PA1-16730 |
| CD3 | Mouse | 1:200 | Invitrogen | 14-0030-82 |
| **Secondary antibodies or conjugated** |  |  |  |  |
| Alexa 488 Anti-CD45 | Mouse | 1:400 | Abcam | Ab256254 |
| Alexa 647 Anti-CD68 | Rabbit | 1:400 | Cell signaling | 17846 |
| Anti-Mouse Alexa 488 | Goat | 1:1000 | Invitrogen | A11001 |
| Anti-Rabbit IgG Alexa 568 | Donkey | 1:1000 | Invitrogen | A10042 |
| Anti-Mouse IgG Alexa 647 | Goat | 1:1000 | Cell signaling | 4410S |

**Supplementary Figure 1.**
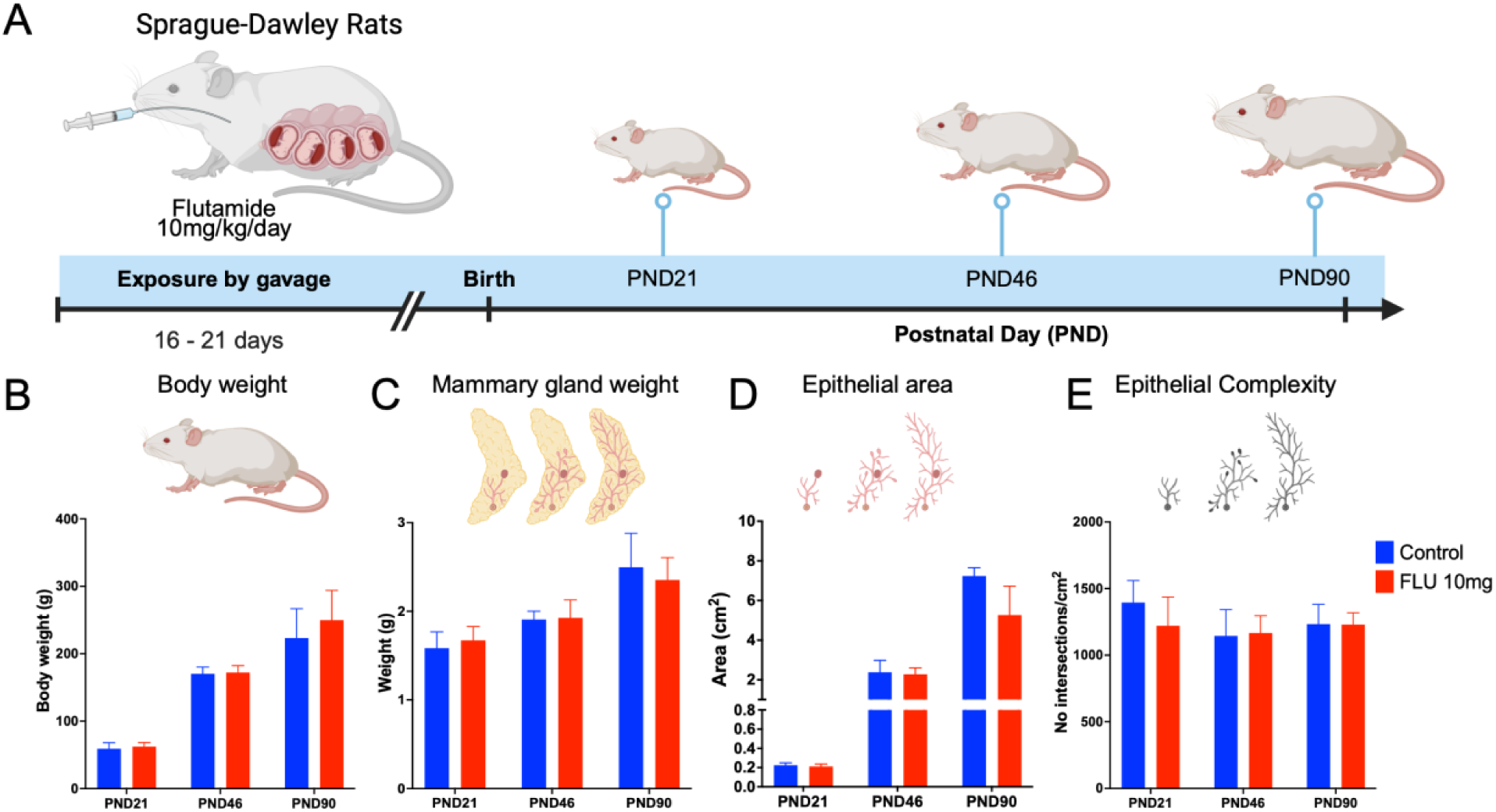
Illustrates the morphological changes in the epithelial mammary gland that occur after in utero exposure to flutamide. **(A)** Representative schematization of *in utero* exposure to flutamide at 10 mg/kg/day. (**B)** Body weight and (**C)** Mammary gland weight at PND21, PND46, and PND90 (*n*=5-7 animals/groups). (**D**) Quantification of total epithelial area and (**E**) epithelial complexity determined by Sholl analysis. Graphs represent the average of the group with SD.

**Figure Supplementary 2.**
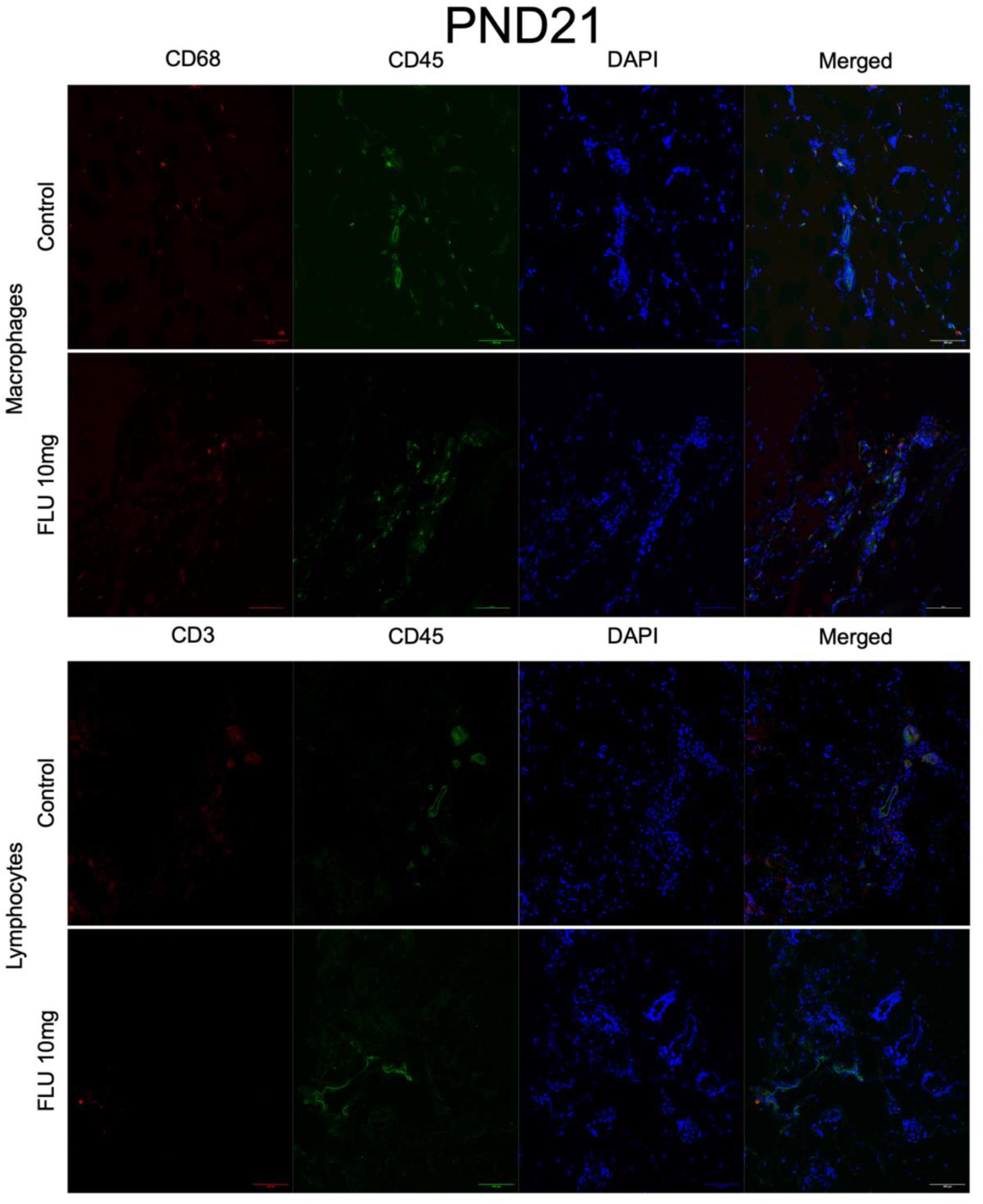
Immune cell quantification at PND21 after FLU exposure in the mammary gland. Immunofluorescence staining using markers CD45 and CD68 for macrophages (**A**) and CD45 and CD3 for lymphocytes (**B**). Magnification 10x, scale bar 300 μm.

**Figure Supplementary 3.**
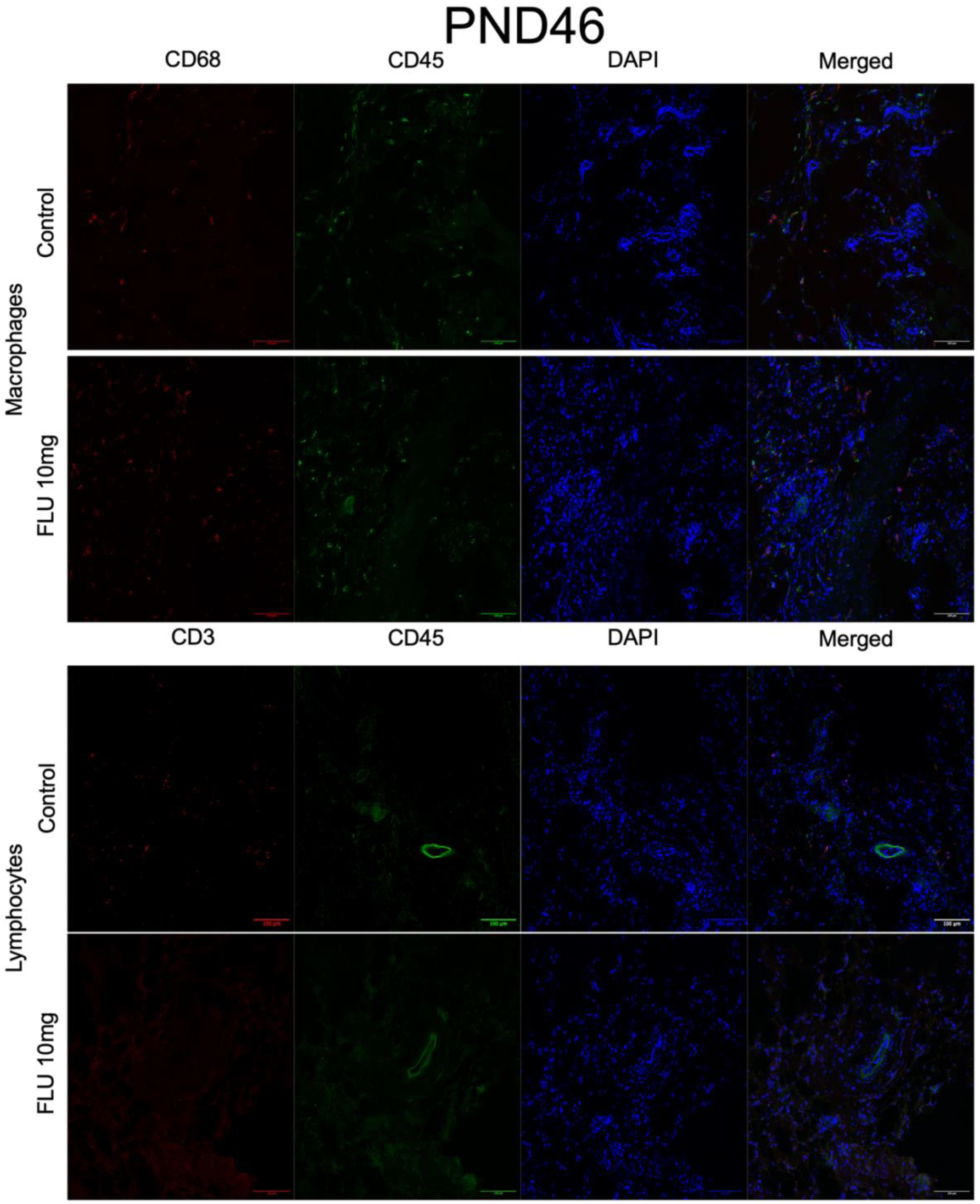
Immune cell quantification at PND46 after FLU exposure in the mammary gland. Immunofluorescence staining using markers CD45 and CD68 for macrophages (**A**) and CD45 and CD3 for lymphocytes (**B**). Magnification 10x, scale bar 300 μm.

**Figure Supplementary 4.**
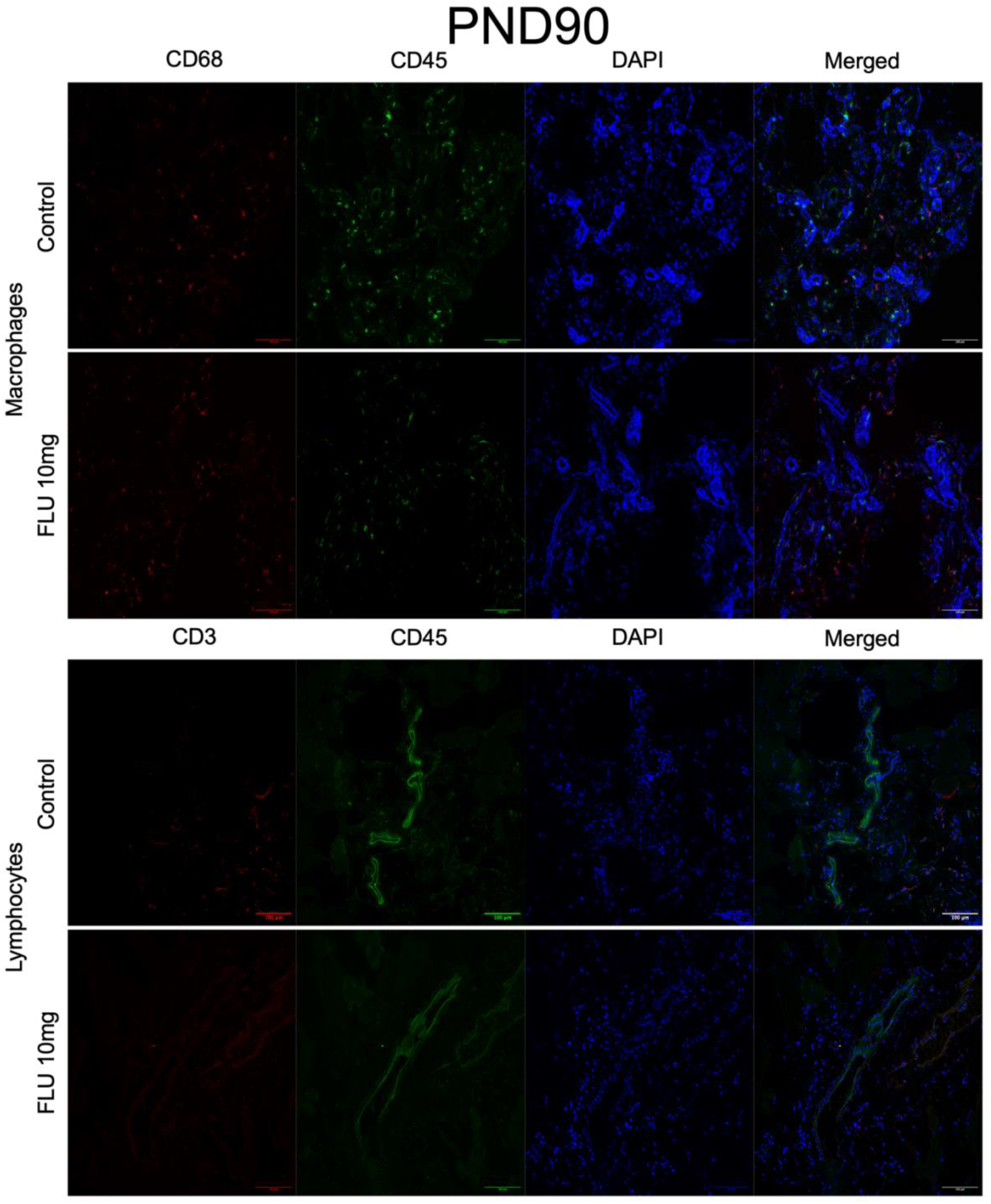
Immune cell quantification at PND90 after FLU exposure in the mammary gland. Immunofluorescence staining using markers CD45 and CD68 for macrophages (**A**) and CD45 and CD3 for lymphocytes (**B**). Magnification 10x, scale bar 300 μm.

